# Spring haul-out behavior of seals in the Bering and Chukchi seas

**DOI:** 10.1101/2022.04.07.487572

**Authors:** Josh M. London, Paul B. Conn, Stacie M. Koslovsky, Erin L. Richmond, Jay M. Ver Hoef, Michael F. Cameron, Justin A. Crawford, Andrew L. Von Duyke, Lori Quakenbush, Peter L. Boveng

## Abstract

Ice-associated seals rely on sea ice for a variety of activities, including pupping, breeding, molting, and resting. In the Arctic, many of these activities occur in spring (April – June) as sea ice begins to melt and retreat northward. Rapid acceleration of climate change in Arctic ecosystems is therefore of concern as the quantity and quality of suitable habitat is forecast to decrease. Improved estimates of seal population abundance are needed to properly monitor the impacts of these changes over time. In this paper, we use hourly percent-dry data from satellite-linked bio-loggers deployed between 2005 and 2021 to quantify the proportion of seals hauled out on ice. This information is needed to accurately estimate abundance from aerial survey counts of ice-associated seals (i.e., to correct for the proportion of animals that are in the water while surveys are conducted). In addition to providing essential data for survey ‘availability’ calculations, our analysis also provides insights into the seasonal timing and environmental factors affecting haul-out behavior by ice-associated seals. We specifically focused on bearded (*Erignathus barbatus*), ribbon (*Histriophoca fasciata*), and spotted seals (*Phoca largha*) in the Bering and Chukchi seas. Because ringed seals (*Phoca (pusa) hispida*) can be out of the water but hidden from view in snow lairs analysis of their ‘availability’ to surveys requires special consideration; therefore,they were not included in this analysis. Using generalized linear mixed pseudo-models to properly account for temporal autocorrelation, we fit models with covariates of interest (e.g., day-of-year, solar hour, age-sex class, wind speed, barometric pressure, temperature, precipitation) to examine their ability to explain variation in haul-out probability. We found evidence for strong diel and within-season patterns in haul-out behavior, as well as strong weather effects (particularly wind and temperature). In general, seals were more likely to haul out on ice in the middle of the day and when wind speed was low and temperatures were higher. Haul-out probability increased through March and April, peaking in May and early June before declining again. The timing and frequency of haul-out events also varied based on species and age-sex class. For ribbon and spotted seals, models with year effects were highly supported, indicating that the timing and magnitude of haul-out behavior varied among years. However, we did not find broad evidence that haul-out timing was linked to annual sea-ice extent. Our analysis emphasizes the importance of accounting for seasonal and temporal variation in haul-out behavior, as well as associated environmental covariates, when interpreting the number of seals counted in aerial surveys.

## NTRODUCTION

Global climate disruption is causing considerable reduction in Arctic sea ice extent, volume, and seasonal presence (Meier et al., 2014; Wang et al., 2018; Kwok, 2018; Overland, 2021). These changes have tangible effects on Arctic organisms, ecosystems, and the people who live in the region (Huntington et al., 2020). Such disruptions are a particular cause of concern for the ice-associated seals that depend on spring and early summer sea ice (March-June) in the Bering and Chukchi seas as a platform for important life history functions, such as pupping, nursing, breeding behavior, and molting (Boveng et al., 2009, 2013; Cameron et al., 2010; Kelly et al., 2010). Limited data and large knowledge gaps complicate predictions about the ultimate effects of changes in sea ice on the behavior, health, abundance, and distribution of these seals. To date, indices of seal health sampled during periods of declining sea ice differ regionally (Crawford, Quakenbush & Citta, 2015; Harwood et al., 2020). Knowledge about evolutionary constraints on the timing of reproductive and molting behavior is generally lacking, so it is difficult to predict how or if ice-associated seal species will adjust to future changes (e.g., by adjusting pupping or molting schedules to earlier dates or different locales). This is further complicated by the spatio-temporal variation in the phenology of these life history events within regions and throughout their full ranges. Additionally, trends in abundance of these species are unknown, so it is difficult to assess the effect, if any, declines in sea-ice habitat have had, or will have, on seal demography.

Statutory requirements (e.g., United States Marine Mammal Protection Act (MMPA), United States Endangered Species Act (ESA)) for timely estimates of population abundance and trend mean improved aerial survey effort is needed for these species. Those survey efforts must also be paired with improved knowledge of haul-out behavior to ensure appropriate survey design, robust methods, and accurate estimates. Several studies have contributed estimates of the distribution and abundance of ice-associated seal species in the Arctic using aerial surveys (e.g., Bengtson et al. (2005), Conn et al. (2014), and Ver Hoef et al. (2014)) and more recent efforts have significantly expanded on previous survey effort. Such abundance studies are conducted over very large areas and estimation of absolute abundance requires making inference about numerous issues affecting the observation of seals on ice. These include availability (only seals on ice are available to be counted), detection probability (observers or automated detection systems may miss some seals on ice), species misclassification, and possible disturbance of seals by aircraft (Ver Hoef et al., 2014; Conn et al., 2014). Refining these inferences will improve the accuracy of abundance estimates and, hopefully, allow credible predictions about the effects of climate disruptions on the abundance and distribution of Arctic seal populations.

How ice-associated seals use sea ice as a haul-out platform varies among species. Ribbon seals (*Histriophoca fasciata*) haul out of the water almost exclusively on sea ice and are mostly pelagic outside the spring pupping, breeding, and molting season (Boveng & Lowry, 2018). Although spotted (*Phoca largha*) and bearded (*Erignathus barbatus*) seals occasionally rest on coastal land, they primarily use sea ice as a haul-out platform during the spring and early summer (Frost & Burns, 2018). Ringed seals (*Phoca (pusa) hispida*) — not included in this study — haul out on sea ice but primarily use snow lairs on sea ice during winter and spring.

The remoteness of the Bering and Chukchi seas means direct scientific observation of seal behavior is impractical. Thus, bio-logging devices are especially useful tools for collecting key information on movement and haul-out behavior for these species. Bio-logging records of time spent out of the water provide valuable data for identifying covariates that explain variation in haul-out behavior. For instance,Von Duyke et al. (2020) used satellite-linked bio-loggers to corroborate seasonal changes between diurnal and nocturnal haul-out behavior of ringed seals previously described by Kelly and Quakenbush (1990) using VHF radio tags and direct observation. Bengtson et al. (2005) documented a higher propensity for ringed seal basking near solar noon, as did Ver Hoef et al. (2014) in an analysis of bearded, ribbon, and spotted seals using much larger sample sizes. Olnes et al. (2020) showed that the proportion of time bearded seals spent hauled out progressively increased through spring and summer, and Ver Hoef et al. (2014) found haul-out probabilities increased gradually starting in March and peaked in May and June for bearded, ribbon, and spotted seals. Such analyses have not been limited to the Arctic. In the Antarctic, Bengtson and Cameron (2004) relied on bio-logging data to demonstrate greater haul-out propensity in juvenile crabeater seals (*Lobodon carcinophaga*) than adults, with highest probabilities in February and at times close to solar noon.

Knowledge of haul-out patterns is not only important for understanding natural history and ecology, but also for developing “availability” correction factors for aerial surveys. Specifically, researchers need to know the fraction of seals hauled out (versus in the water) when aerial surveys are conducted. Studies estimating availability correction factors for seals typically use logistic regression-style analyses to estimate the time-specific probability of being hauled out based on ‘wet/dry’ data relayed by bio-loggers. In these models, haul-out probabilities were expressed as a function of predictive covariates, such as time-of-day, day-of-year, sex, age class, and environmental conditions (e.g., Reder et al. (2003), Bengtson & Cameron (2004), Bengtson et al. (2005), Udevitz et al. (2009), Lonergan et al. (2013), Ver Hoef et al. (2014), Southwell et al. (2008), and Niemi et al. (2023)). However, sample sizes have often been insufficient to permit strong inference about demographic and/or seasonal variation in haul-out probabilities. For instance, Bengtson and Cameron’s (2004) study included 5 adult and 2 juvenile crabeater seals, while Bengtson et al.’s (2005) study was based on 6 ringed seals in the Chukchi and Beaufort seas. These studies were often further limited by logistical constraints on fieldwork and the attachment duration or operational life of bio-loggers. In this study, we addressed some of these limitations by deploying small bio-loggers designed for longer-term attachment on rear flippers of a subset of the study individuals. These devices are designed to collect data through the molt period (when those adhered to the hair would fall off) and, in some situations, provide multiple years of data.

In this study, we used 16 years of bio-logging data to investigate the haul-out behavior of bearded, ribbon, and spotted seals in the Bering and Chukchi seas. Our goals were threefold. First, we wished to establish baseline estimates for the chronology of haul-out behavior in the critical spring season for each species across different age and sex classes. Second, we sought to refine estimates of haul-out availability corrections for aerial surveys in order to improve estimates of seal abundance. Previously estimated availability correction factors (e.g., Bengtson et al. (2005), Conn et al. (2014), and Ver Hoef et al. (2014)) accounted for variables such as the time-of-day and day-of-year, but did not investigate the impact of weather variables. Such variables have been shown to influence walrus haul-out behavior (Udevitz et al., 2009) and we expect weather conditions to also influence seal haul-out behavior and including them within the model framework will benefit our estimates of seal availability during aerial surveys. Third, we aimed to assess the annual variability in haul-out timing and possible linkage to changes in the extent of seasonal sea ice between 2005 and 2021. Our work extends the scope of previous haul-out analyses, includes the influence of weather variability, and investigates the potential impact of changing sea-ice extent on the behavior of these species.

## METHODS

### Data collection

For this study we used haul-out behavior data and location estimates from bio-loggers deployed on bearded, ribbon, and spotted seals in the Bering, Chukchi, and western Beaufort seas by multiple organizations as part of collaborative investigations from 2005 through 2021. Seals were captured using nets and bio-loggers were attached during studies based in coastal communities or on research ships. Ship-based capture events occurred during spring near the southern ice edge in the Bering Sea between 2005 and 2018. Land-based capture events occurred between 2005 and 2020 from May to October, generally between the Alaska coastal communities of Scammon Bay in the Bering Sea, Utqiaġ vik in the Chukchi Sea, and Nuiqsut in the Beaufort Sea (Supplemental Material, S1). Data from additional deployments along the Kamchatka peninsula in the western Bering Sea are also included. We refer readers to Figure 1 and the primary literature for detailed capture and bio-logger attachment methods (see also Supplemental Material, S1).

**Figure 1.**
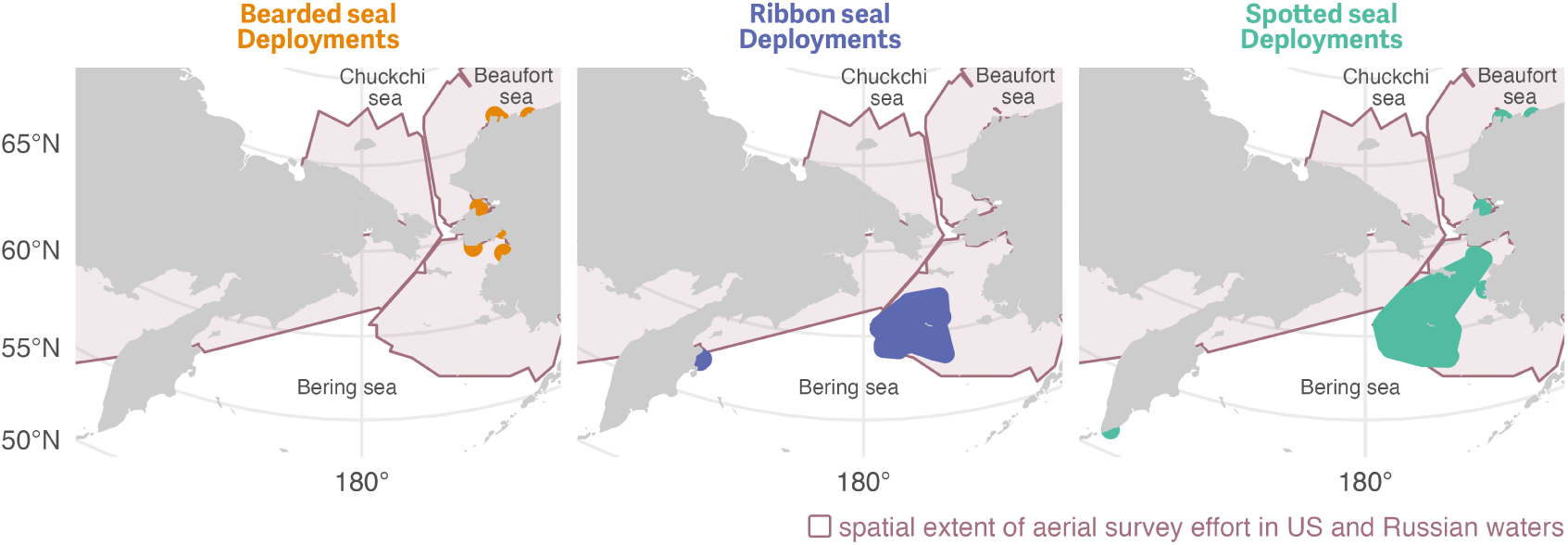
Initial bio-logger deployment areas for bearded, ribbon, and spotted seals in the study between 2005 and 2020 across multiple regions within the Bering, Chukchi, and western Beaufort seas. Solid regions shown for each species are minimum concave polygons buffered by 60 km for enhanced visibility. The larger, shaded region indicates the spatial extent of aerial survey effort to date in the Bering, Chukchi, and Beaufort seas. Deployments were initiated across a range of months but only haul-out data from 1 March to 15 July were included in the analysis. See Figure 3 for the spatial distribution of observed data used in the study and Supplemental Materials S1 for additional deployment details.

We created a subset of haul-out behavior data from 250 bio-loggers deployed on 35 bearded, 110 ribbon, and 105 spotted seals to include only records from 1 March to 15 July between 2005 and 2021. Bio-loggers were of the ‘SPLASH’ or ‘SPOT’ family of tags developed by Wildlife Computers (Redmond, Washington, USA). Deployments consisted of either a single ‘SPLASH’ device, a single ‘SPOT’ device, or both types. Devices were either adhered to the hair on the seal or attached through the rear flipper inter-digital webbing. The use of bio-loggers adhered to the back or head provides some benefits over flipper mounted devices (e.g. increased satellite transmittal rates, locations at sea) but these fall off during the following annual molt, which, depending on deployment date, limits the duration of haul-out data they provide especially during the focus months of our study.

Additionally, bio-loggers attached to the head or dorsal region are often dry while the seal is floating at the surface, inducing a slight positive bias in the hourly percent-dry values reported by the bio-logger. For this study, in cases where both bio-logger types were deployed, we preferred hourly percent-dry observations from the flipper tag. All data were transmitted by the deployed instruments via the Argos satellite network and location data were either derived from Argos transmissions or transmitted FastLoc GPS data.

Sex and age class (non-dependent *young-of-the-year*, sexually immature *subadults*, and mature *adults*) were estimated at the time of deployment by various combinations of length, claw growth ridges (McLaren, 1958), and pelage characteristics. Seals determined to be less than one year were classified as young-of-the-year. For those bio-loggers deployed on young-of-the-year and transmitting into the next year, the age class was advanced to subadult on 1 March of the following year – the assumed anniversary of their birth. Subadults are those seals likely greater than one year of age but less than four years. Adults are individuals that are likely older than four years. Table 1 provides a summary of these deployments and data received from them.

**Table 1.** Summary of bio-logger data by seal species and age classification from 1 March to 15 July 2005-2021. Seal hours represents the sum of hourly observations across all seals used in the analysis. Because young-of-the-year are advanced to subadult on 1 March of the following year, some individual seals are represented in both columns in this table

| Species | Sex | Age Class |  |  |
| --- | --- | --- | --- | --- |
|  |  | Adult | Subadult | Young-of-the-Year |
| Bearded seal | F | 1 ( 1,776 seal hours) | 16 (21,648 seal hours) |  |
| Bearded seal | M | 2 ( 2,108 seal hours) | 16 (17,232 seal hours) |  |
| Ribbon seal | F | 33 (35,128 seal hours) | 18 (15,984 seal hours) | 13 ( 3,734 seal hours) |
| Ribbon seal | M | 24 (27,465 seal hours) | 19 (13,046 seal hours) | 9 ( 4,228 seal hours) |
| Spotted seal | F | 23 (21,588 seal hours) | 21 (19,559 seal hours) | 11 (13,417 seal hours) |
| Spotted seal | M | 20 (31,793 seal hours) | 21 (17,210 seal hours) | 12 (11,285 seal hours) |

Haul-out behavior data were recorded in a manner standard across Wildlife Computers bio-loggers and transmitted via the Argos satellite network as hourly percent-dry timelines. For each hour of a day, the wet/dry sensor was polled by the tag firmware every few seconds and the percent of the hour in the dry state was calculated (Figure 2). On board the bio-logger, hourly percent-dry calculations were rounded to the nearest 10% inclusive of 0% and 100% along with additional values at 3% and 98%. This compression resulted in additional data transmission as each message consisted of two complete 24-hour records. Memory capacity allowed caching of percent-dry records for several weeks or months and each message was transmitted several times to ensure reception at the satellite. Bio-loggers were deployed and programmed in a manner to maximize data transmission during the spring pupping and molting period, though hourly percent-dry data were not always successfully transmitted. This is due to a variety of factors including satellite coverage, tag availability (e.g., tags mounted to the rear flipper often do not transmit while at sea), tag performance, duty cycling, and atmospheric interference. Fortunately, missing records do not substantially bias inference about haul-out probabilities (Conn et al., 2012).

**Figure 2.**
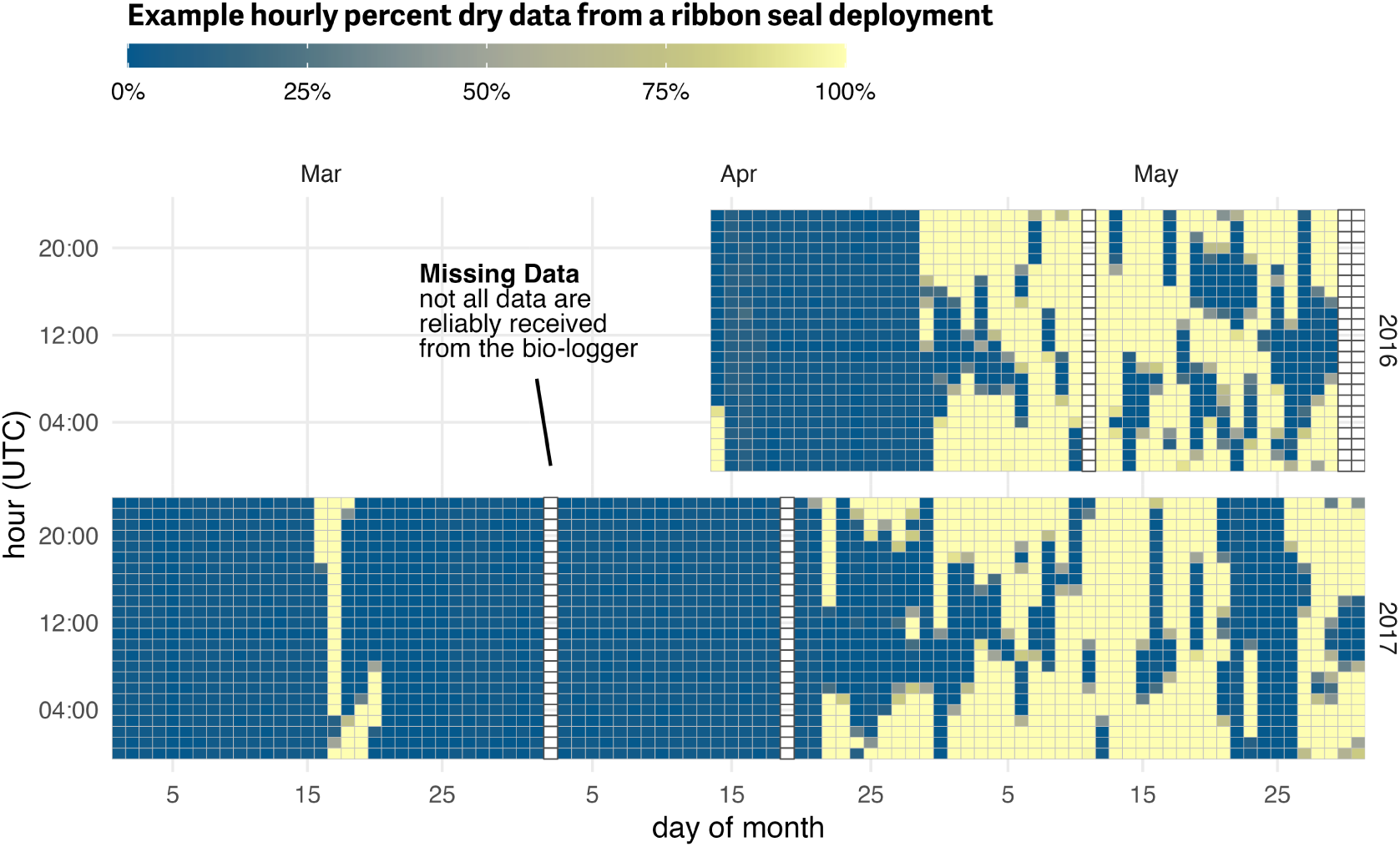
Example percent-dry actogram from bio-logger data. Haul-out behavior observations recorded by a bio-logger deployed on a ribbon seal over two years during the months of March, April, and May. This actogram plot represents the transmitted hourly percent-dry values. Not all data during a deployment are reliably transmitted from the bio-logger and because data are transmitted in 24-hour chunks occasional missing days are present in the record. Here, missing data that were not successfully received from the device are represented as empty rectangles.

Tags that fell off due to molt, attachment failure, or seal mortality and remained on ice or land may have continued to send data to satellites; i.e., a detached bio-logger that is dry (either on ice or land) will record and transmit data suggesting the seal is hauled out. Therefore, end times of each deployment were identified by examining bio-logger locations, percent-dry records, and dive behavior (if available) to determine when bio-loggers ceased providing data consistent with seal behavior. For example, a data record that ended with several consecutive days (∼10+ days) of 100% dry observations and with locations indicating the tag was on land were truncated to the final stretch of 100% dry observations. The vast majority of deployments ended with the device detaching in the water and the deployment end date was obvious. There is no perfect algorithm for identifying deployment end dates and each deployment in question was considered separately. While not perfect, we are confident our reliance on expert opinion and examination of multiple data streams provided the best option. Data outside of the deployment start and end times were discarded prior to analysis.

Of key interest in this study was the relationship between haul-out behavior and weather covariates that vary with time and seal location. The use of modern bio-loggers that record and transmit behavioral data while simultaneously providing location estimates was key to this objective. We explored the use of a continuous-time correlated random walk (Johnson et al., 2008) movement model to predict locations at specific times. However, the sparse nature of data from some bio-loggers, especially those mounted to the rear flipper, resulted in poor modeling performance or convergence issues. For this study, we calculated a weighted average daily location where the inverse of the estimated Argos or FastLoc GPS location error was used for the weight. Each Argos location estimate was assigned an error radius based on either the categorical location quality ( *3* = 250 m, *2* = 500 m, *1* = 1500 m, *0* = 2500 m (Lopez et al., 2013); we chose 2500 m for location classes *A* and *B*) or, when available, the estimated error radius from the Argos Kalman filter algorithm. Location estimates from FastLoc GPS were all assigned an error radius of 50 m. On days when haul-out observations were present but location data were missing we used the seal’s last calculated weighted average daily location; days when the location intersected with land were removed from the seal’s record. We recognize that bearded and spotted seals haul out on land. However, assessing the relationship between haul-out behavior and weather covariates and seals’ availability for aerial surveys on land was outside the scope of this study. Additionally, any daily locations on land were likely more reflective of coordinate averaging and measurement error, rather than actual use of coastal features.

### Explanatory variables

In addition to sex and age class, we analyzed variables that might help explain variation in haul-out probabilities. These included day-of-year (for seasonal effects) and local solar hour (for diurnal effects). We calculated local solar hour using the {solaR} package (Perpiñán, 2012) within the R statistical environment (R Core Team, 2021) based on the weighted daily average locations. We also linked the weighted average daily locations to weather values from the North American Regional Reanalysis (NARR) model produced by the National Centers for Environmental Prediction (Mesinger et al., 2006). The NARR model assimilates observational data to produce a long-term picture of weather over North America and portions of the surrounding seas. Weather variables are made available across the region 8 times daily. For this study, NARR weather values were limited to the extent of our study area over the Bering, Chukchi, and Beaufort seas at 3-hr intervals based on the original grid resolution of 32 km (1024 km^2^). The following weather variables are known to affect haul-out behavior in other Arctic pinnipeds (Reder et al., 2003; Udevitz et al., 2009; Perry, Stenson & Buren, 2017) and were interpolated and assigned to daily seal locations using a bilinear method: 1) air temperature at 2 m above the Earth’s surface, 2) wind consisting of northerly and easterly vector components converted to wind speed using the Euclidean norm, 3) barometric pressure at sea level, and 4) precipitation (Table 2).

**Table 2.** Explanatory covariates used in analyses of binary haul-out records for bearded, spotted, and ribbon seals. Note that we also considered select interactions (see article text) between these primary covariates. For instance, wind chill was represented by the interaction temperature:wind.

| Covariate | Type | Description |
| --- | --- | --- |
| Age-sex class | Categorical | young-of-the-year, subadult, adult male and adult female |
| Hour | Continuous;<br>Fourier basis | local solar hour using 6 variables of a Fourier-series basis |
| Day | Continuous | linear, quadratic, and cubic effects of day-of-year |
| Precip | Continuous | convective precipitation kg/m<sup>2</sup> (NARR) |
| Pressure | Continuous | atmospheric pressure at sea level (kPa) (NARR) |
| Temp | Continuous | air temperature (C) at 2m above the earth's surface (NARR) |
| Wind | Continuous | northerly and easterly vector components for wind (NARR) converted into a single wind speed via the Euclidean norm |
| Northing | Continuous | latitude divided by the mean latitude across all locations (for bearded seals only) |
| Year | Continuous | For the set of models examining inter-annual variation in sea-ice use, we fitted models with the addition of year by day-of-year interactions. |

For all seal species, we considered the following variables when modeling the hourly haul-out behavior: day-of-year, solar hour, temperature, wind speed, barometric pressure, precipitation, and wind chill (represented by a *wind:temperature* interaction (Udevitz et al., 2009)). Ribbon and spotted seal models included age-sex class and interactions between day-of-year and age-sex class, but we omitted these from bearded seal models due to poor representation of age-sex classes (Table 1). Bearded seal models included a latitudinal effect (and an interaction with day-of-year), because bearded seals occupy a substantial range during the spring and we were interested in possible differences in the timing of haul-out behavior along a latitudinal gradient. We omitted the latitudinal effect from ribbon and spotted seal models because, during the spring, these species are most prevalent near the southern ice edge in the Bering Sea (Conn et al., 2014).

### Haul-out modeling

Haul-out records for seals are often characterized by sequential hours spent hauled out on ice alternating with long periods in the water (Figure 2). Commonly used statistical models for binary data (e.g. logistic regression) assume independence among responses, an assumption that is clearly violated if hourly responses are modeled. Any analysis that ignores temporal autocorrelation in responses will thus have overstated precision (Betts et al., 2006).

To properly account for temporal dependence and to take advantage of computational efficiency, we used generalized linear mixed pseudo-models (GLMPMs; Ver Hoef, London & Boveng (2010)) to model variation in haul-out behavior as a function of (1) covariate predictors, (2) temporally autocorrelated random effects, and (3) individual random effects representing heterogeneity in individual behavior. We used the glmmLDTS package (Ver Hoef, London & Boveng, 2010) to implement GLMPMs. We explored two different model formulations for our data and we fit separate models to bearded, ribbon, and spotted seal data sets as we expected differing behavior by species. Separate models for each species were also needed because a single, very large data set proved computationally intractable. In our first model formulation and for each species, we fitted a year-independent model that predicted average haul-out behavior as a function of demographic, weather, seasonal, and diurnal effects. Second, for ribbon and spotted seals (which had considerably more data than bearded seals), we fitted models that included all the effects from the first model, but also permitted annual variation in haul-out timing. This second set of models was used to examine whether haul-out patterns varied by year and to determine the annual timing of apparent peaks in haul-out behavior. For both models, we assumed an hourly Bernoulli response (a binary 0-1 response dependent upon whether the tag was greater than 50% dry for a given hour) where the linear predictor was modeled on the logit scale. This is consistent with previous approaches (London et al., 2012; Ver Hoef et al., 2014) and only 6.995% of our observations fell between 10% and 90% hourly percent-dry.

We followed Ver Hoef et al. (2014) in using linear, quadratic, and cubic effects of day-of-year to represent temporal changes in behavior during the season. However, unlike previous models for harbor seals (London et al., 2012) and ice-associated seals (Ver Hoef et al., 2014), which treated hour-of-day as a 24-level categorical variable to capture diurnal cycles, we adopted a continuous formulation based on Fourier series that provides a flexible model while preserving the inherent circularity needed for time-of-day effects (i.e., hour 0 should be equal to hour 24). It also represents hour-of-day with 6 parameters, which is a considerable reduction when compared to a 24-parameter variable. According to this approach, we used the following specification for hour-of-day effects:

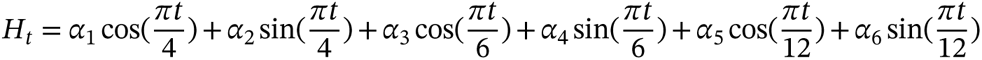

where 𝐻_𝑡_ gives the effect for solar hour 𝑡 and 𝛼_𝑖_ are estimated parameters (regression coefficients). For the second set of models examining inter-annual variation in sea-ice use, we fitted models with year by day-of-year interactions. However, in this case we only included *year:day* and *year:day^2^*, omitting the main effects of year as well as *year:day^3^* interactions because models with the latter effects were numerically unstable. However, the modeled interactions were sufficient to allow shifts in haul-out distribution, as one can show mathematically that a simple horizontal shift in timing of haul-out distributions does not affect the main effects or cubic terms in a polynomial regression model. Bearded seals were not included in this examination of inter-annual variation because of limited data across many years in the study.

A typical model fitting exercise would also include a model selection process. However, AIC (and similar criteria) is not suitable when using pseudo-likelihoods, because pseudo-data generated in the model fitting process (Ver Hoef, London & Boveng, 2010) differ between models (Ten Eyck & Cavanaugh, 2018). After fitting GLMPM models, we instead used “type III” 𝐹-tests to calculate 𝑝-values (Ver Hoef, London & Boveng, 2010) to evaluate model performance and important terms. We also produced predictions of haul-out behavior as a function of three influential predictors (e.g. solar hour, day-of-year, age-sex). Weather covariates for these predictions were based on daily or hourly smoothed weather covariate values across the study region. Such predictions were then used to develop haul-out probability surfaces, explore conditional effects of weather covariates, and determine annual peaks in haul-out activity. The timing of peak haul-out behavior was further used to regress against the annual maximum sea-ice extent in the study region. Predictions before 15 March and after 30 June were not included in visualizations or other evaluations to avoid spurious model predictions at the edge of the data range.

Visualizing the marginal or conditional effect of an individual weather covariate (where all other weather covariates are being held at mean values) on haul-out probability was difficult in this analysis because of the collinearity between covariates as well as the spatial and temporal variation across such a large region. The relationship of each weather covariate with haul-out probability, averaged over the other weather conditions, was more variable than model coefficients would imply. That said, important insights can be gained from plots of marginal effects. To create these plots, we predicted haul-out probability across the full range of each weather covariate while fixing hour of the day at local solar noon and day-of-year at 15 May. For the other weather covariate values, we chose not to use a fixed mean value because we expect weather to vary within day over the season (e.g. the temperature at solar noon will gradually increase from March through June). To account for this, we fit a simple generalized additive model for each weather covariate with smooth terms for day-of-year and solar hour. We used predicted values from the generalized additive model in lieu of holding other weather covariates at a fixed mean value which would not capture seasonal change. The visualizations also include vertical lines representing 95% confidence intervals around the predicted haul-out probability to better communicate the variation in model uncertainty.

We also assessed whether the annual variation in maximum spring sea-ice extent in the Bering Sea influenced the seasonal peak of seal haul-out probability. In particular, we used sea-ice concentration data from the Nimbus-7 SMMR and DMSP SSM/I-SSMIS Passive Microwave Dataset, Version 1 (Cavalieri et al., 1996) to calculate maximum sea-ice extent. All sea-ice concentration grid cells (25 km^2^) in the study area with greater than 15% concentration were counted daily to get the total sea ice extent for each day between 15 February and 15 July across all years. Maximum spring sea-ice extent was simply the largest daily count of grid cells with greater than 15% concentration for each year. A separate regression model, built on the results of the haul-out model, was used to evaluate the relationship between the annual computed peak haul-out day (as the response) with the maximum sea-ice extent (as the predictor).

## RESULTS

Figure 3 shows the spatial distribution of weighted locations with available haul-out behavior data used for analysis of each species across the study area. Figure 4 shows the temporal distribution of all haul-out data across the study season for each species. Observations of ribbon and spotted seals were concentrated in the months of May and June due to the timing of deployment (April and May) and the timing of molt (May and June). During molt, seals (and their attached bio-loggers) spend more time out of the water and more data are transmitted. Molt timing also impacts when many deployments end as any bio-loggers adhered to the hair will fall off. Relative to the other species in the study, there were fewer deployments of bio-loggers on bearded seals. This resulted in fewer data observations overall and noticeably lower in numbers May and June. The majority of deployments on bearded seals occurred in August and September and, by May, bio-loggers had either fallen off or their batteries were depleted. Therefore, observations for bearded seals were concentrated in March (Figure 4).

**Figure 3.**
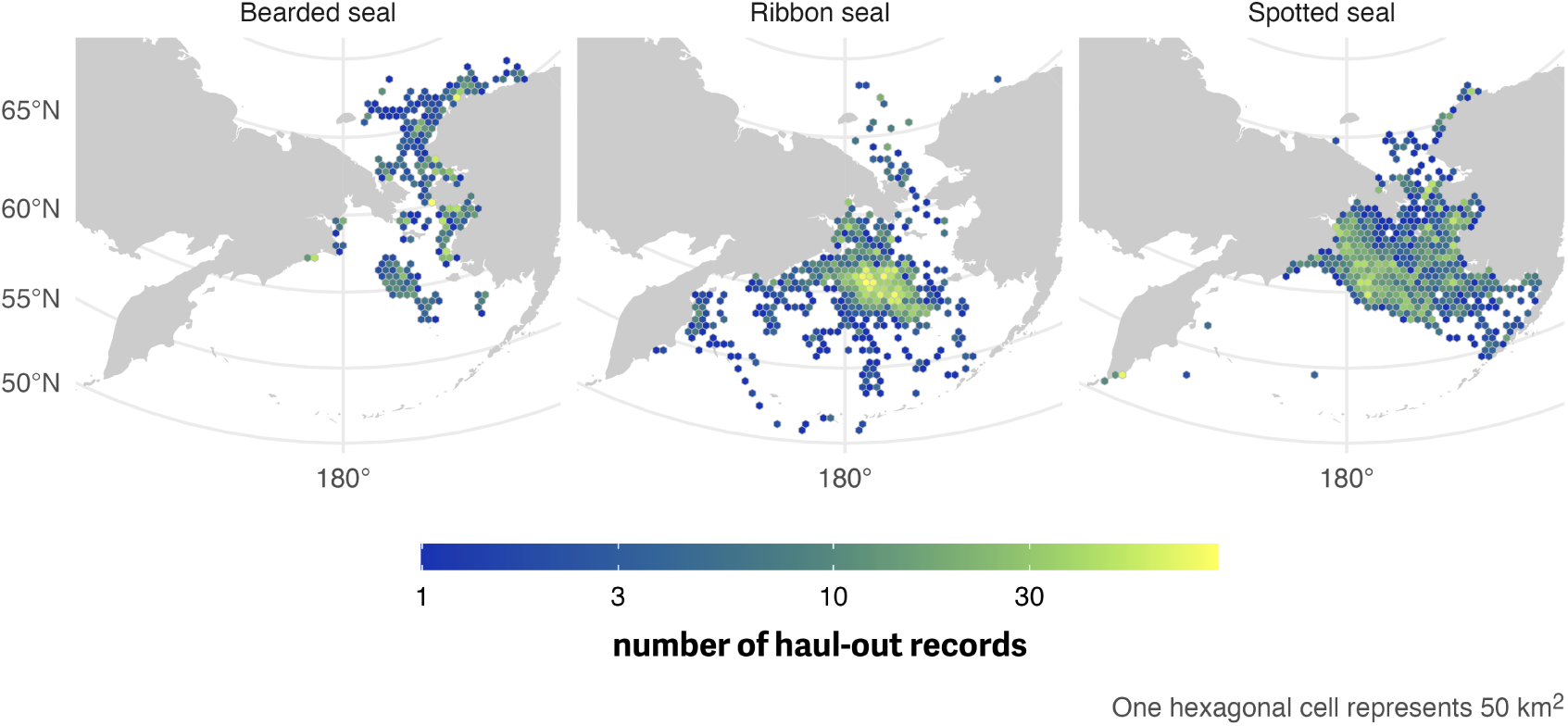
Spatial distribution of haul-out records from 1 March through July 15 for bearded, ribbon, and spotted seal. Linking location estimates with haul-out records in space and time allows for inclusion of weather covariates in the final model. For this visualization, data were collated across all years between 2005 and 2020 and each hexagonal cell represents an area of 50 km^2^

**Figure 4.**
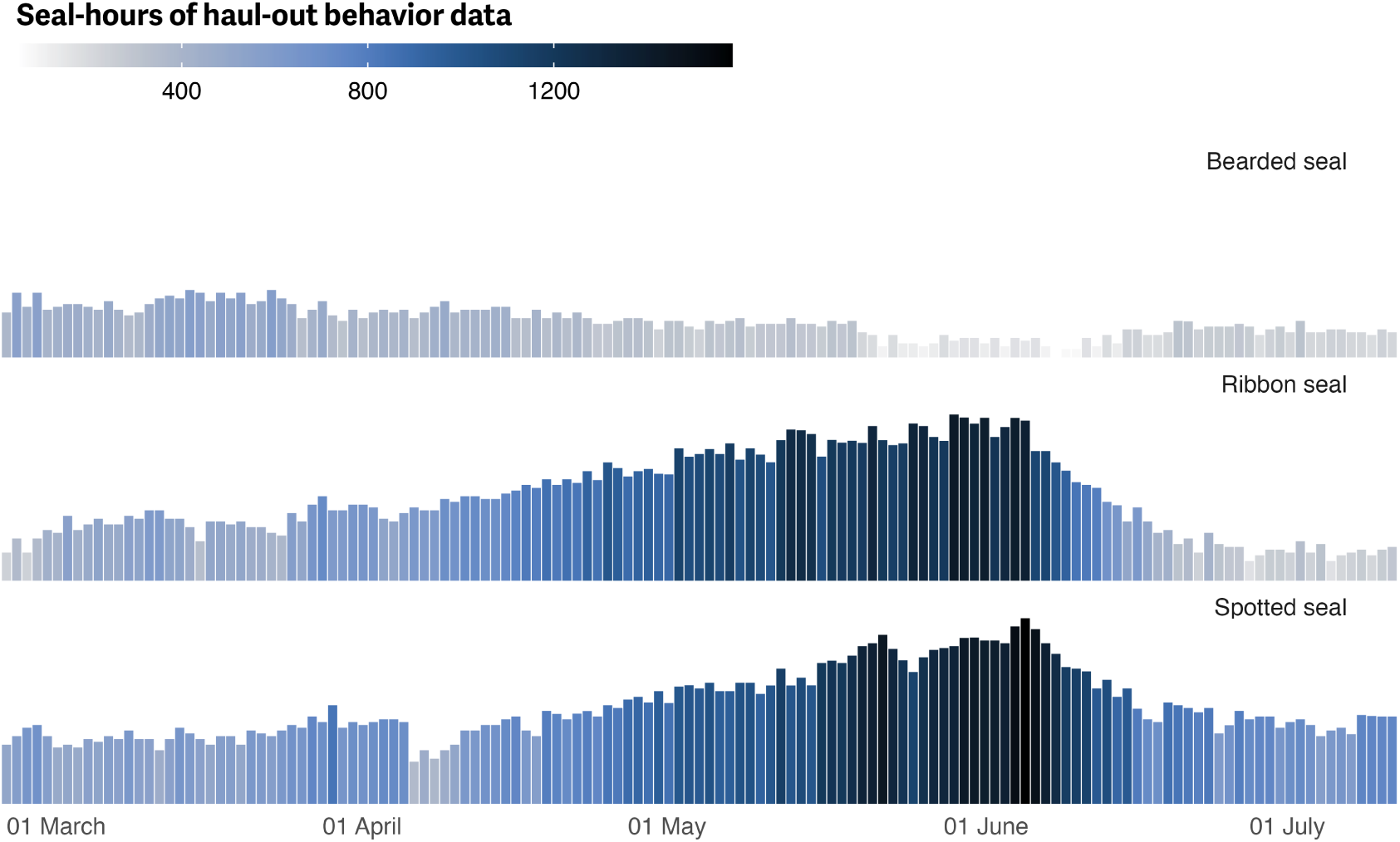
Seasonal distribution of haul-out behavior observations by species. Distribution of hourly percent-dry bio-logger data from 1 March to 15 July for each species. Data are grouped by day-of-year and presented in seal-hours collated across all years between 2005 and 2020. Seal-hours of data are represented as a color gradient and bar height. The higher amount of data from May and June in ribbon and spotted seals coincides with peak molting when seals (and their attached bio-loggers) are more likely hauled out. Additionally, many bio-logger deployments started in April and May. The overall reduced quantity of observations from bearded seals is reflective of the lower number of bio-logger deployments in the study and the fact that the majority of deployments on bearded seals occurred in August and September; therefore most bio-loggers deployed had exhausted their batteries.

Across all three seal species, generally, models omitting year effects suggested that day-of-year, solar hour, age-sex class, temperature, and wind substantially influenced haul-out behavior of all three species, with 𝐹 tests producing 𝑝-values less than 0.05 for variables embodying these effects and/or their interactions. Haul-out probabilities typically increased throughout March and April, reaching a peak in May and early June before declining again. Diurnal patterns were present, with maximum haul-out behavior centered around local solar noon.

### Bearded Seals

Age and sex class were not included in the model for bearded seals due to our lower sample size for adult and young-of-year age classes. As such, results are shown for all ages (Figure 5; see also S1). Additionally, after approximately 9 months, 7 devices deployed on the rear flipper of bearded seals reported implausible hourly percent-dry data (100% dry for several weeks but indicative of movement and increasing transmission rates (see Boveng & Cameron (2013))). All data after the first instance of unrealistic values were censored from this analysis. In addition to a peak around local solar noon, the bearded seal model predicted additional haul-out activity around local midnight. In concert with the lower magnitude of haul-out probability, bearded seal haul-out behavior was also more protracted throughout the spring season compared to ribbon (Figure 7) and spotted seals (Figure 9). Overall, bearded seals were less likely to haul out and had a bi-modal distribution of haul-out probability across the day.

**Figure 5.**
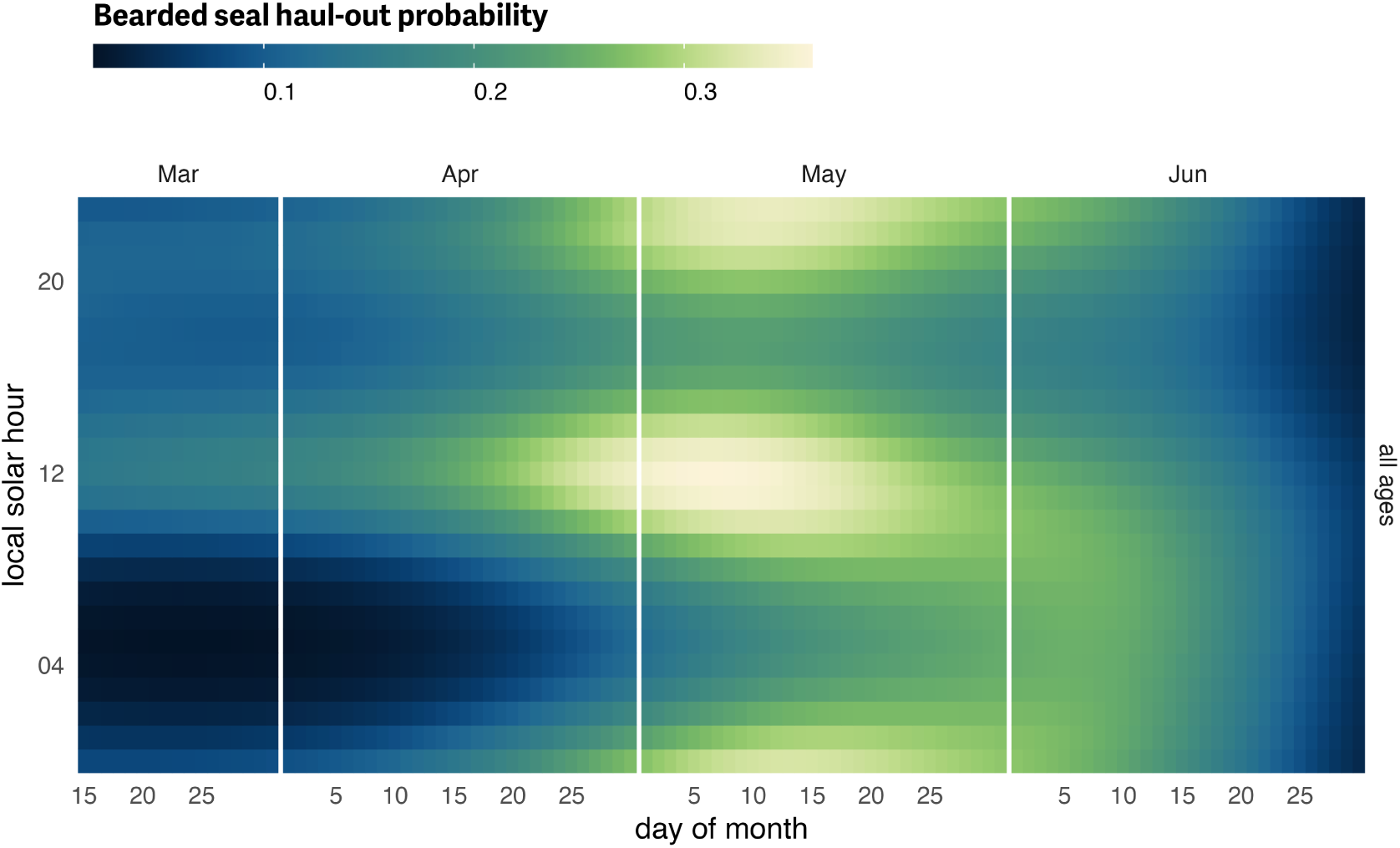
Bearded seal predicted haul-out probability. Predicted hourly haul-out probability of bearded seals (all ages and sex classes) from 15 March to 30 June for all age and sex classes combined. Predictions in July and before 15 March were not included to avoid spurious model predictions at the edge of the data range. Weather covariate values in the prediction were based on a simple generalized additive model for each weather covariate with smooth terms for day-of-year and solar hour to account for anticipated variability within a day over the season. Note, our model predicts lower haul-out probability for bearded seals overall compared to ribbon and spotted seals and the color scales are not directly comparable.

When exploring the influence of weather, bearded seal haul-out probability was strongly affected by wind (𝐹_1,42728_ = 130.468; 𝑝 = <0.001) and temperature (𝐹_1,42728_ = 19.5; 𝑝 = <0.001) with much higher haul-out probability during periods of higher temperatures and low wind speeds (Figure 6). Not surprisingly, wind chill (𝐹_1,42728_ = 14.54; 𝑝 = <0.001) was also important. Barometric pressure (𝐹_1,42728_ = 7.779; 𝑝 = 0.005) was also significant factor although less apparent (Figure 6). Any effect of precipitation was not a significant influence on haul-out probability (𝐹_1,42728_ = 0.519; 𝑝 = 0.471).

**Figure 6.**
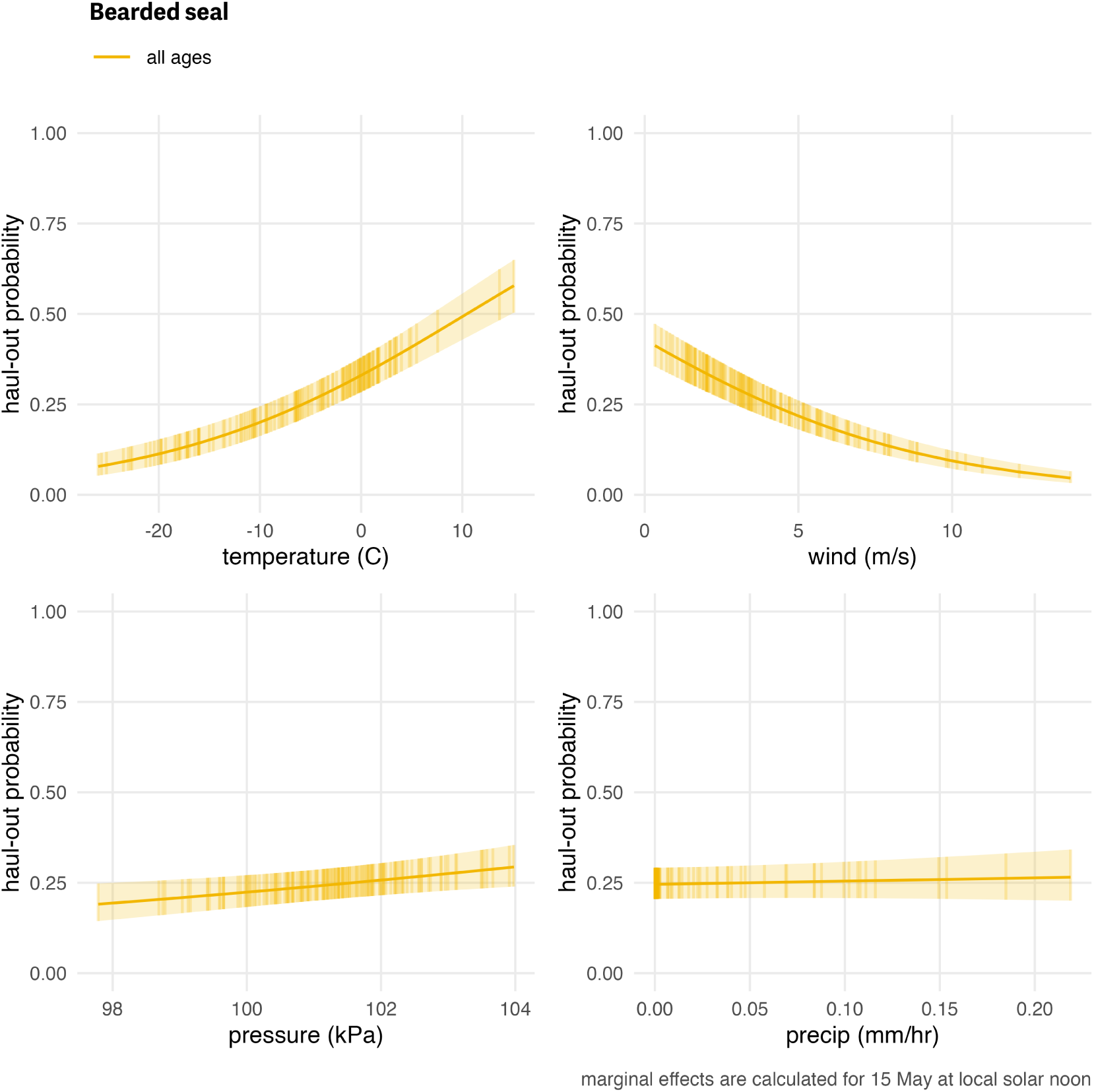
Influence of weather covariates on bearded seal haul-out probability. Marginal effects of temperature, wind, barometric pressure, and precipitation on the predicted haul-out probability of bearded seals combined across all age and sex classifications. Hour of the day was fixed at local solar noon and day-of-year fixed at 15 May. The other weather covariate values were predicted from a simple generalized additive model for each weather covariate with smooth terms for day-of-year and solar hour to account for anticipated variability within a day over the season. While the marginal effect of the covariate is continuous, points and vertical lines representing the 95% confidence interval around the predicted haul-out probability are shown only at observed weather values to give an indication of how the observations are distributed across the range of weather values.

### Ribbon Seals

Ribbon seals exhibited a pattern of gradually increasing haul-out probability in April that peaks in late May for subadults and in early June for adults (Figure 7; see also S2). There is an apparent seasonal progression with subadults hauling out earlier in the season followed by adult males and, then, adult females. Haul-out behavior was clearly centered around local solar noon and expanded to other hours later in the spring as seals entered their molting period. Subadults showed an earlier start and more intense haul-out activity in April and May. The young-of-the-year records began after weaning and the model predictions seemed to indicate development of in-water activities (e.g. swimming, foraging) in May. Adult females had a more protracted haul-out season compared to males, and more time was spent hauled out in June compared to adult males and subadults.

The haul-out probability for ribbon seals was mostly influenced by temperature (𝐹_1,99540_ = 6.87; 𝑝 = 0.009) and wind (𝐹_1,99540_ = 49.314; 𝑝 = <0.001) with barometric pressure (𝐹_1,99540_ = 3.446; 𝑝 = 0.063) having a milder impact. Ribbon seals were more likely to haul out when temperatures were relatively warm and less likely to haul out at higher winds and lower pressure values (Figure 8). Neither wind chill (𝐹_1,99540_ = 1.83; 𝑝 = 0.176) nor precipitation (𝐹_1,99540_ = 0; 𝑝 = 0.989) were a significant influence on haul-out probability. Compared with bearded seals, the effect of weather covariates on the predicted haul-out probability for ribbon seals was less striking. Because our ribbon seal model included age and sex class, we can visualize the different influences of weather covariates on those classes and see that subadults differ from adult males and females (Figure 8).

**Figure 7.**
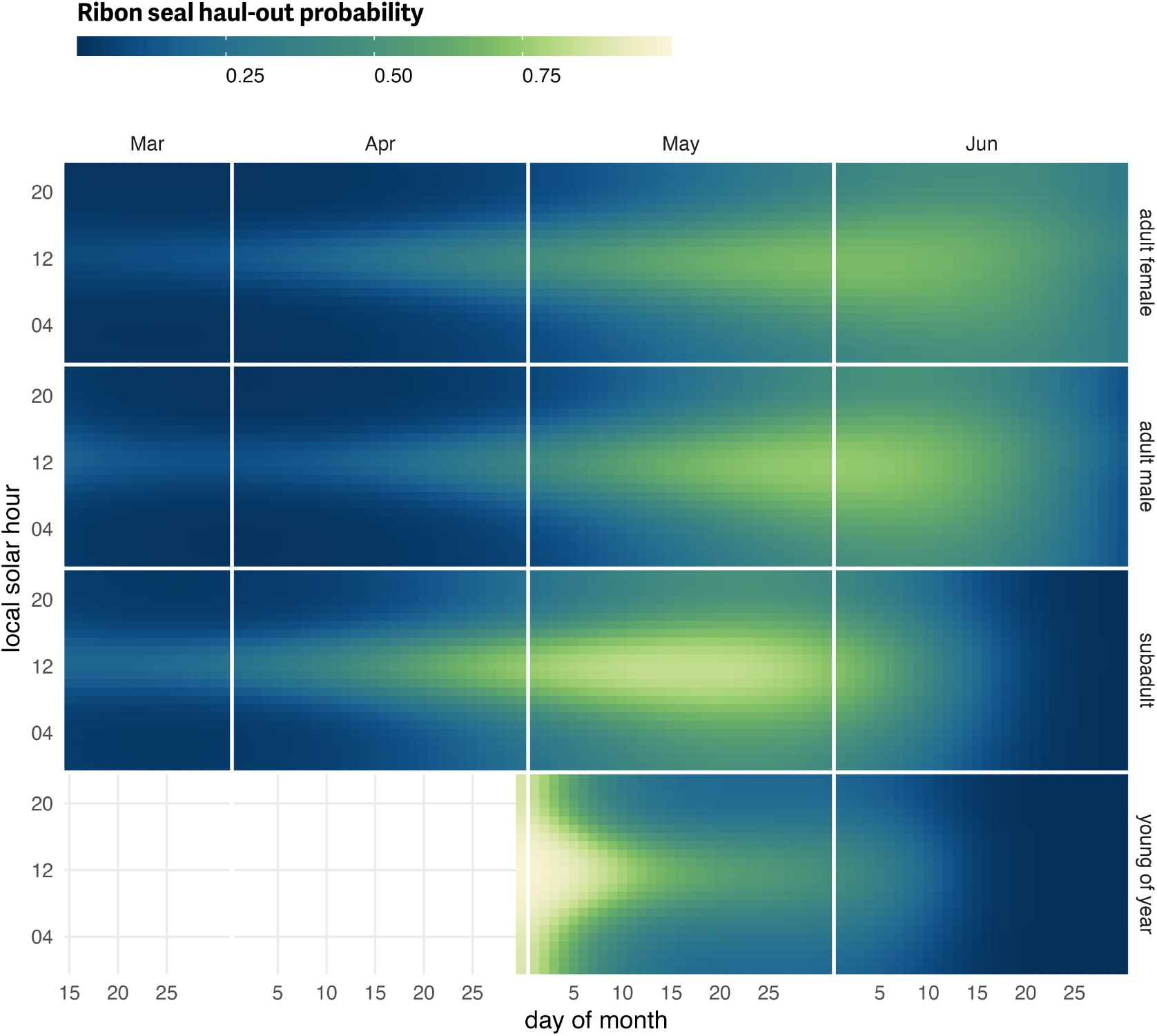
Ribbon seal predicted haul-out probability. Predicted hourly haul-out probability of ribbon seals from 15 March to 30 June for each age and sex class used in the model. Weather covariate values in the prediction were based on a simple generalized additive model for each weather covariate with smooth terms for day-of-year and solar hour to account for anticipated variability within a day over the season. Predictions for young of the year show their transition from newly weaned pups resting on the ice to more in-water activities. Predictions in July and before 15 March were not included to avoid spurious model predictions at the edge of the data range.

**Figure 8.**
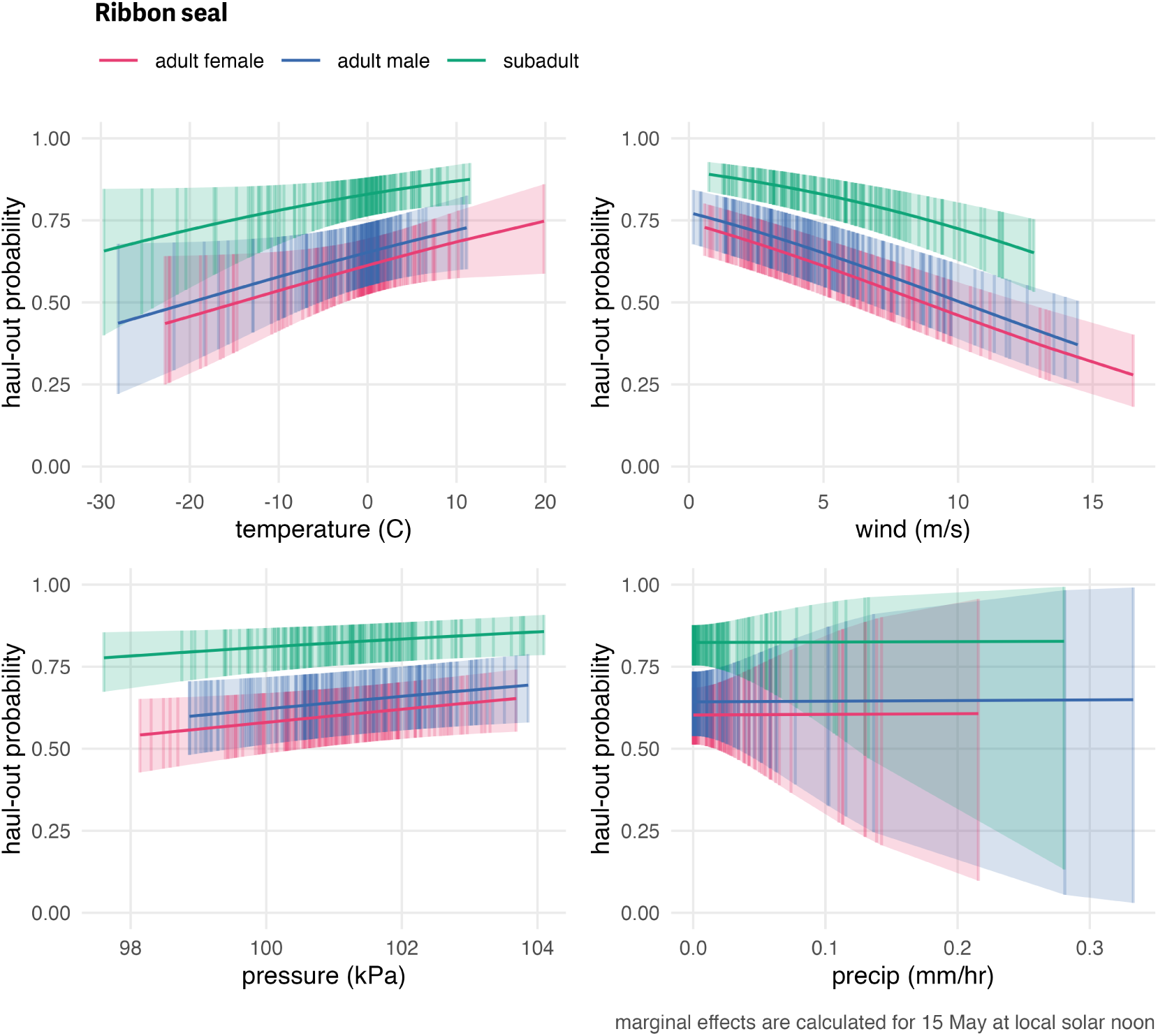
Influence of weather covariates on ribbon seal haul-out probability. Marginal effects of temperature, wind, barometric pressure, and precipitation on the predicted haul-out probability of ribbon seals within each age and sex classifications. Hour of the day was fixed at local solar noon and day-of-year fixed at 15 May. The other weather covariate values were predicted from a simple generalized additive model for each weather covariate with smooth terms for day-of-year and solar hour to account for anticipated variability within a day over the season. While the marginal effect of the covariate is continuous, points and vertical lines representing the 95% confidence interval around the predicted haul-out probability are shown only at observed weather values to give an indication of how the observations are distributed across the range of weather values.

**Figure 9.**
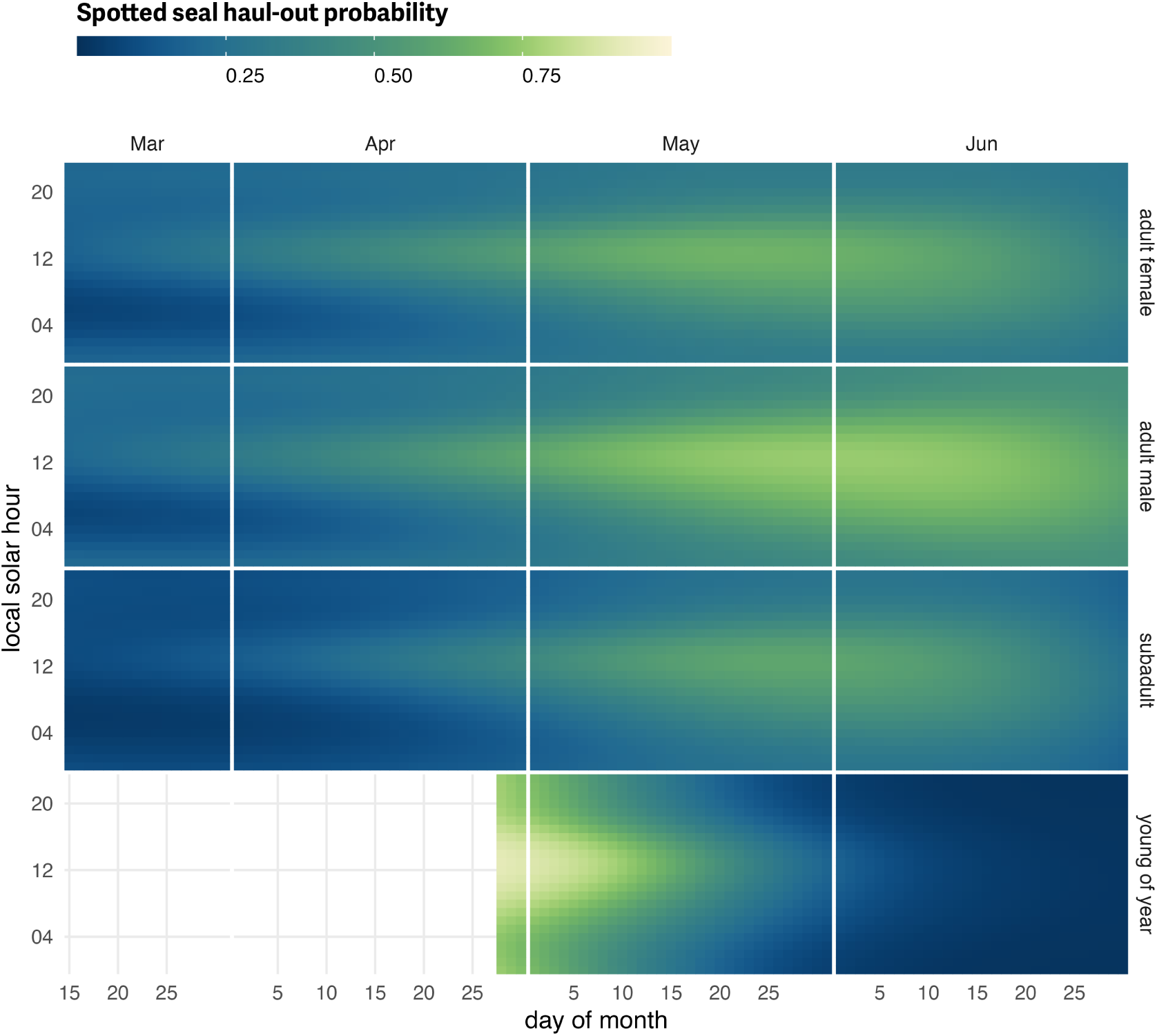
Spotted seal predicted haul-out probability. Predicted hourly haul-out probability of spotted seals from 1 March to 30 June for each age and sex class used in the model. Weather covariate values in the prediction were based on a simple generalized additive model for each weather covariate with smooth terms for day-of-year and solar hour to account for anticipated variability within a day over the season. As with ribbon seals, predictions for young of the year show their transition from newly weaned pups resting on the ice to more in-water activities. Predictions in July and before 15 March were not included to avoid spurious model predictions at the edge of the data range.

### Spotted Seals

Compared to ribbon seals, spotted seals showed a longer spring haul-out season that was less intensely centered on solar noon (Figure 9; see also S3). Adults of both sexes spent considerable time in April, May and June hauled out. Adult spotted seal males had a more protracted haul-out season compared to females, and more time out of the water in June (Figure 9). This likely reflects the triad behavior in spotted seals when suitor males haul out with nursing females. As with ribbon seals, the young-of-the-year records began after weaning and the model predictions reflected development of in-water activities (e.g. swimming, foraging) in May.

Spotted seal haul-out behavior was less affected by the weather covariates compared to ribbon and bearded seals but their influence on the model was still significant in some cases. Temperature (𝐹_1,114807_ = 5.462; 𝑝 = 0.019), wind (𝐹_1,114807_ = 46.954; 𝑝 = <0.001), and barometric pressure (𝐹_1,114807_ = 10.214; 𝑝 = 0.001) were all significant. Spotted seals were more likely to be on the ice when temperatures were relatively warm and less likely to haul out at higher wind speeds. Wind chill (𝐹_1,114807_ = 0.559; 𝑝 = 0.455) and precipitation (𝐹_1,114807_ = 0.763; 𝑝 = 0.382) were not as influential as the other covariates. Differences in the magnitude of response between the age-sex classes were present and consistent across each of the weather covariates (Figure 10). There was a consistent ranking of adult males being the most likely to haul out, followed by adult females, and, then, subadults. This differs from ribbon seals which showed more overlap between adult males and adult females and that subadults were most likely to haul out across the presented range of weather covariates.

**Figure 10.**
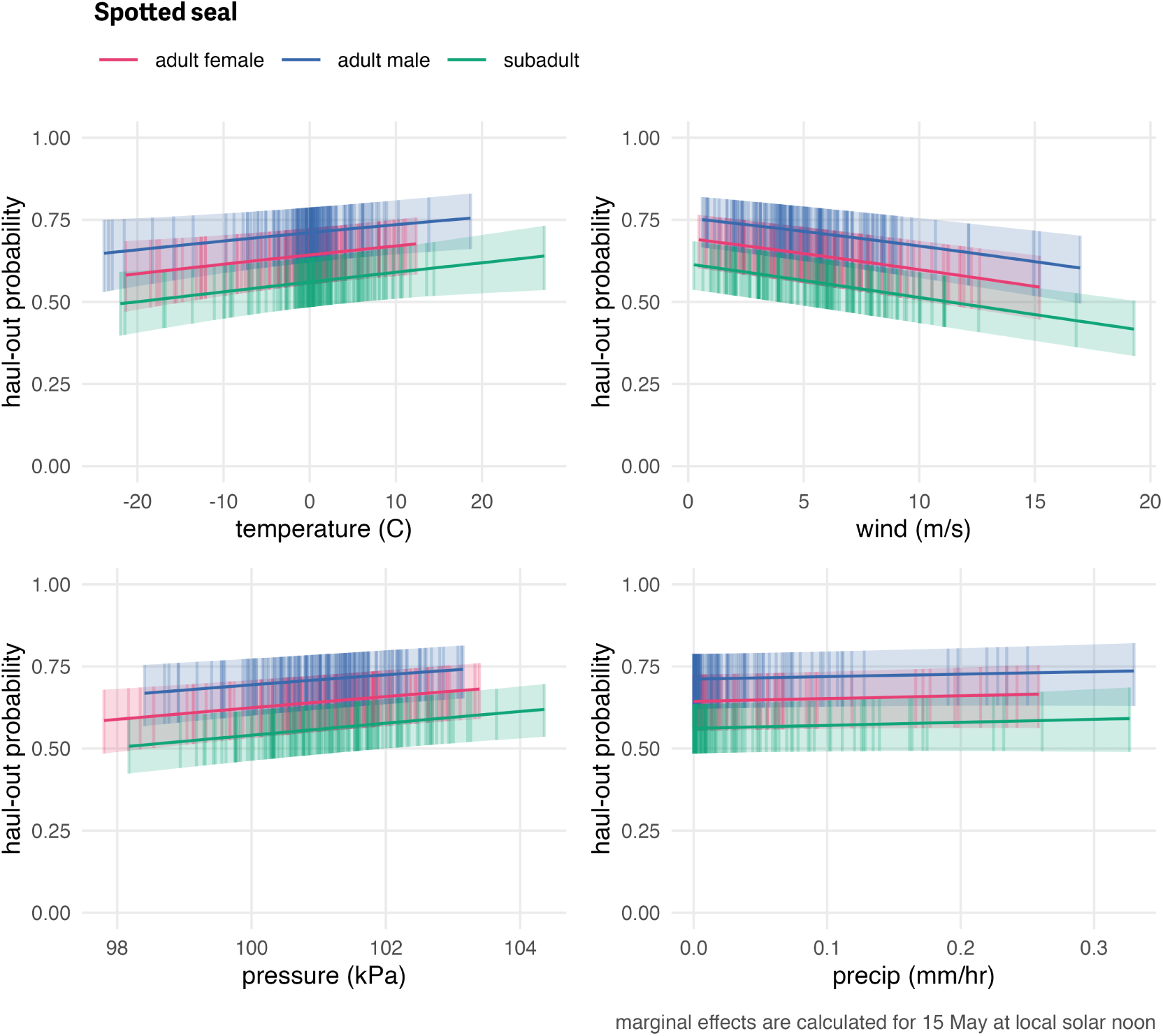
Influence of weather covariates on spotted seal haul-out probability. Marginal effects of temperature, wind, barometric pressure, and precipitation on the predicted haul-out probability of spotted seals within each age and sex classification. Hour of the day was fixed at local solar noon and day-of-year fixed at 15 May. The other weather covariate values were predicted from a simple generalized additive model for each weather covariate with smooth terms for day-of-year and solar hour to account for anticipated variability within a day over the season.

### Annual variation in haul-out timing

The second set of models, which included annual variation in haul-out patterns, uncovered significant contributions for linear and quadratic interactions between day and year for only spotted seals (day:year, 𝐹_15,114762_ = 4.345; 𝑝 = <0.001; day^2^:year, 𝐹_15,114762_ = 5.827; 𝑝 = <0.001). Ribbon seals showed no significant contribution for interactions between day and year (day:year, 𝐹_10,99510_ = 0.516; 𝑝 = 0.880; day^2^:year, 𝐹_10,99510_ = 0.549; 𝑝 = 0.856). Predicted distributions of haul-out activity were largely unimodal, but varied some among and within years with respect to the timing and magnitude of haul-out peaks (Figure 11). It is important to note that predicted variation in annual haul-out patterns likely reflected both process error and sampling variability. While we did remove any years

**Figure 11.**
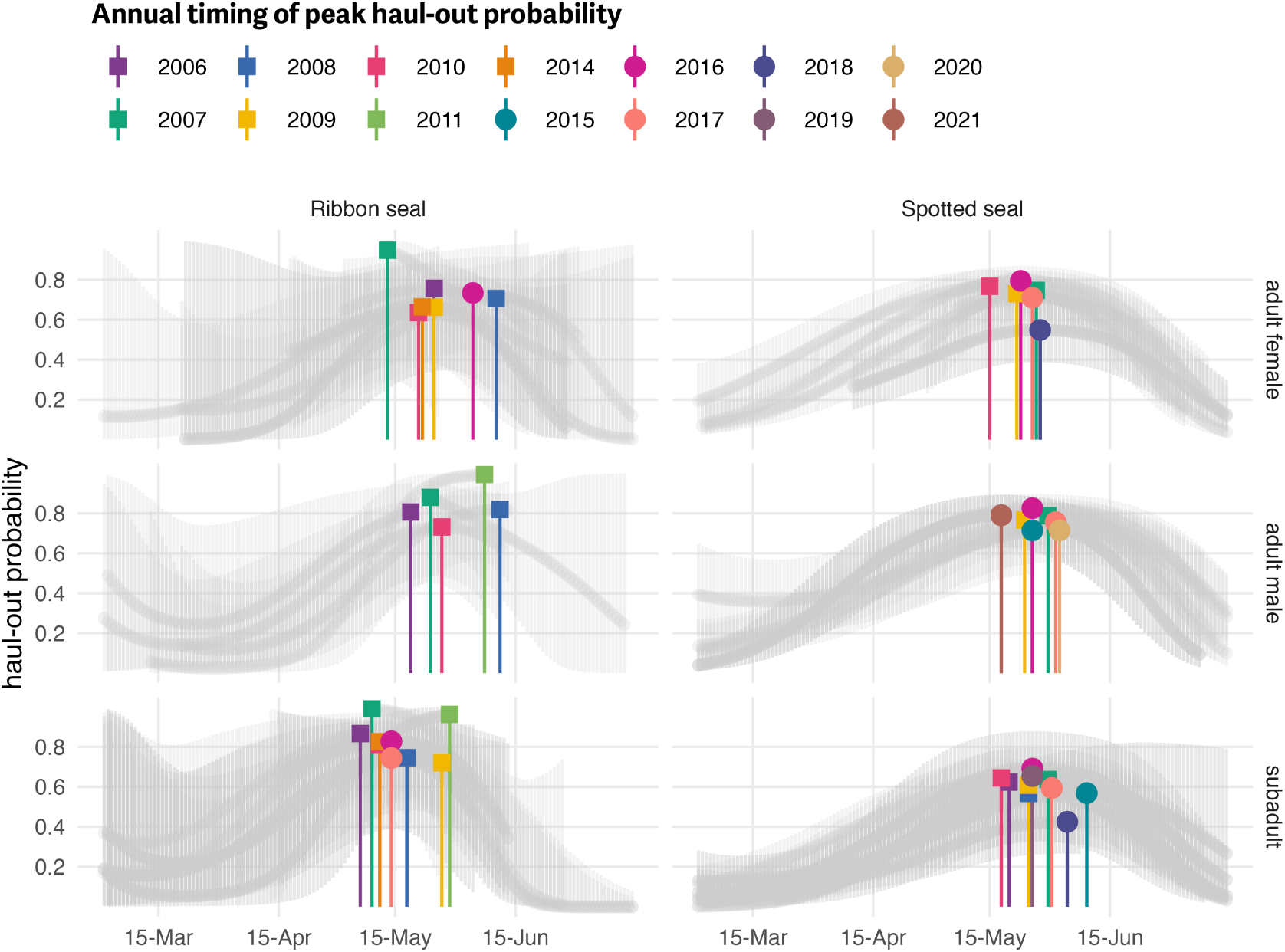
Annual variability in the timing of peak haul-out probability for ribbon and spotted seals. Spring haul-out probability (colored markers) for ribbon and spotted seals across 14 years are shown. The full seasonal range of annual predictions along with uncertainty are presented as grey lines. Covariates were fixed at local solar noon and weather covariate values were predicted from a simple generalized additive model for each weather covariate with smooth terms for day-of-year and solar hour to account for anticipated variability within a day over the season. Only those groups (age:sex + year) that included observations from more than one seal are shown. Additionally, any groups where data were only available after 1 June or before 1 May are not included.

While the marginal effect of the covariate is continuous, points and vertical lines representing the 95% confidence interval around the predicted haul-out probability are shown only at observed weather values to give an indication of how the observations are distributed across the range of weather values. where only one deployment in a species + age:sex group was present, there were still some years where the pattern shown was informed by a small number of individuals that may not represent population-level patterns.

The annual timing of peak haul-out probability for ribbon seals and adult male spotted seals appeared to have no relationship with the amount of yearly maximum sea-ice extent in the study area. For ribbon seals and adult male spotted seals, 𝑝-values were substantially larger than 0.05 (ribbon seal adult females; 𝑅^2^ = 0.004, 𝑝 = 0.896; ribbon seal adult males: 𝑅^2^ = 0.059, 𝑝 = 0.693; ribbon seal subadults: 𝑅^2^ = 0.007, 𝑝 = 0.828; spotted seals adult males: 𝑅^2^ = 0.003, 𝑝 = 0.903). Adult female and subadult spotted seals both showed a negative trend (peak haul-out occurs later in years with less sea ice) with a significant relationship for adult female spotted seals (spotted seal adult female: 𝑅^2^ = 0.663, 𝑝 = 0.049; spotted seal subadults: 𝑅^2^ = 0.384, 𝑝 = 0.056).

## DISCUSSION

We modeled data from bio-loggers deployed on bearded, ribbon, and spotted seals to examine factors affecting their haul-out behavior on sea ice in the Bering and Chukchi seas. Our analysis shows all three species of seal haul out progressively more through the spring and peak near mid-May to early June before declining again. This pattern aligns well with what has been previously documented qualitatively (Boveng et al., 2009; Cameron et al., 2010; Boveng & Cameron, 2013) and confirms our haul-out data are likely quantifying population-level behavioral patterns. Beginning in spring, seals preferentially haul out on ice shortly after solar noon, which allows seals to maximize absorption of solar radiation thought to facilitate the molting process (Feltz & Fay, 1966). Interestingly, bearded seals appear to have two peaks in haul-out activity within a day, one shortly after solar noon, and one centered near solar midnight. This, of course, could be an artifact of our limited sample size for bearded seal deployments across all age classes. However, a similar bi-modal pattern has been seen in ringed seals (Von Duyke et al., 2020) and suggests that bearded and ringed seals may be operating under different constraints than ribbon and spotted seals. Bearded and ringed seals are distributed across higher latitudes higher latitudes that experience extended daylight hours during spring which may allow more flexibility in alternating resting and foraging events. Other factors such as predation by polar bears (which is rare for ribbon and spotted seals in the Bering Sea) may also explain differing haul-out patterns. The change in haul-out behavior during the season was less pronounced in bearded seals compared to ribbon and spotted seals. This aligns with findings from Thometz et al. (Thometz et al., 2021) who observed a mean molting period of 119±2 days and a relatively stable resting metabolic rate for bearded seals during that time. While ribbon seals were not considered in that study, spotted and ringed seals underwent molt periods of just 33±4 and 28±6 days and had increased resting metabolic rates.

Unlike previous analyses of seal haul-out behavior in spotted and ribbon seals (e.g. (Ver Hoef, London & Boveng, 2010), (Conn et al., 2014)), we also investigated the influence of sex-age class on haul-out probabilities of both species (but not for bearded seals because of low sample size). Field identification of age class can be inexact, particularly when differentiating subadults from adults. In the case of ribbon seals, subadults often have less distinct ribbons than adults. Spotted seal pelage cannot be used to reliably discern adults from subadults. Despite these challenges, we feel the age classifications used in this analysis are useful in testing for age-related effects on haul-out behavior. Although ribbon and spotted seals exhibited a unimodal diel haul-out pattern generally centered around local solar noon, there were key differences across species, age, and sex that match our understanding from natural history descriptions of their ecological behavior. Spotted seals are known to form triads during the breeding season where a female and dependent pup are accompanied on the ice by a suitor male (Frost & Burns, 2018). The male waits for the female to wean the pup and enter estrus, and fends off any other potential suitor males. Triad formation results in both males and females spending a large portion of the day hauled out on ice and a protracted spring haul-out season for both sexes. While females are still nursing the pup and not yet in estrus they may be less inclined to interrupt their haul out and enter the water where the suitor male would attempt mating. We see this reflected in the predicted haul-out patterns, with both males and females exhibiting a broad distribution of time out of the water throughout the solar day and the season. Ribbon seals are not known to form triads and our model predicts a progression of increased haul-out behavior with females starting earlier in the season than males. Notably, female ribbon seals spend a large portion of the day in the water during the pupping period, aligning with the hypothesis that ribbon seal females continue foraging while nursing. Subadult ribbon and spotted seals begin elevated haul-out behavior earlier in the spring and follow a pattern seen in other phocids where yearlings and subadults molt first followed by adult females and males (Thompson & Rothery, 1987; Kirkman et al., 2003; Reder et al., 2003).

We also investigated the influence of weather on haul-out probabilities, including wind speed, temperature, barometric pressure, precipitation, and wind chill. These have been investigated for walruses (e.g. Udevitz et al. (2009)) and a few select studies of ice-associated seals (Perry, Stenson & Buren, 2017; Hamilton & Evans, 2018). In our study, ribbon seal haul-out behavior seemed to be the most influenced by weather, with wind, temperature, and barometric pressure all being important components of the model. Spotted seals were most affected by wind and barometric pressure. For bearded seals, the model indicated wind and temperature had the greatest impact. In general, and as might be expected, seals were more likely to haul-out when daily temperatures were warmer, winds speeds were lower, barometric pressure was higher, and precipitation was lower. Those weather conditions are general indicators of increased solar radiation and lower convective heat loss, both of which provides energetic benefits (see additional discussion in Supplemental Material Exploring Insolation (Solar Radiation) as a Model Covariate regarding the potential use of solar radiation directly). Low winds and precipitation could also enhance predator detection. Our results highlight the importance of incorporating weather covariates when analyzing haul-out behavior and calculating availability corrections for aerial surveys.

Notably missing from our list of haul-out model explanatory variables is any spatial-temporal representation of sea-ice concentration, area, or extent. This may seem counterintuitive when modeling the haul-out behavior of seal species with such a close association to sea ice; seals haul out in the presence of sea ice, and we could assess the local concentration of sea ice during these events (see (Olnes et al., 2020)). This, however, expands the scope of our analysis into the realm of habitat selection and many of our deployments consisted of a single device attached to the rear flipper of the seal which meant we only received locations when seals were hauled out on sea ice, limiting our ability to fully evaluate fine-scale habitat preferences related to sea ice. Insight into how seals use and interact with sea ice during an extended period when the availability and characteristics of sea ice is rapidly changing is important (Breed et al., 2018; Cameron et al., 2018) but ancillary to the focus of this analysis – and, in the end, not possible given key limitations of our data. Additionally, our study was limited to the spring season when seal haul-out behaviors are strongly influenced by pupping, nursing, breeding behavior, and molt. These drivers are likely more influential on haul-out behavior than sea-ice concentration. Crawford et al. (2019) compared haul-out probability models for ringed seals and found those that only included season (and not sea-ice concentration) were the most parsimonious. For all of these reasons, we have elected not to use sea-ice concentration *as a predictor for haul-out probability* in the present study.

Our models detected small deviations in the timing and magnitude of annual peaks in haul-out behavior for ribbon and spotted seals. The timing of peak haul-out activity appears to fall within a relatively narrow time window of 3-4 weeks in late May and early June. This consistency across 15 years is likely a reflection of the relationship between a critical photoperiod and the timing of important life history stages (Temte, 1994; Bronson, 2009). However, along with a critical photoperiod, ribbon and spotted seals are dependent upon the presence of sea ice for pupping and molt. We did not find large support in our models for a relationship between the timing of peaks in haul-out behavior and the amount of yearly maximum sea ice. This could indicate that, while the extent of spring sea ice in the Bering Sea varied widely during our study period, seals were still able to locate sea ice and haul out. Only a small portion of our data was from 2018 – 2019, years of extreme low spring ice extent in the Bering Sea that appeared to impact body condition of ribbon and spotted seals (Boveng et al., 2020), so we may currently lack sufficient contrast in ice extent to characterize an effect on the timing of peak haul-out probability. We should expect, however, that some minimal threshold in the spatial extent or density of sea ice will have a meaningful impact on the timing of peak haul-out behavior — if there is no sea-ice, seals will not haul out or they will be forced to use terrestrial haul-out sites which were not part of the evolution of their normal behaviors. Additionally, while from an ecological perspective the haul-out behavior appears consistent, the interannual differences in timing and magnitude are large enough to have important ramifications on calculations of abundance and trend. Those ramifications will only be exacerbated if climate variability amplifies interannual differences.

Previous attempts to estimate the abundance of phocid seals from aerial survey data in the Bering and Chukchi seas (e.g. Bengtson et al. (2005), Conn et al. (2014), Ver Hoef et al. (2014)) have used estimated haul-out probabilities to correct for the proportion of animals that are in the water and thus unavailable to be counted. Although several of these studies allowed haul-out probabilities to vary by day-of-year and time-of-day, they have not accounted for variability among years, weather conditions, or in the age-sex class of the sample. In this paper, we have shown that there can be considerable differences in the haul-out probability of seals on ice based on these factors and subsequent analyses have shown the potential for considerable bias in abundance estimates if such covariates are unaccounted for (Conn & Trukhanova, 2023). We recommend that future abundance analyses employ availability models that account for them. For instance, it is relatively straightforward to obtain weather reanalysis products for times and locations that are surveyed and to construct a relevant correction factor based on predictions of GLMPMs. The most challenging element in developing availability correction factors is with annual variability. It can be difficult to get a sufficient sample size to estimate year-specific correction factors, particularly because research teams would likely need to deploy bio-loggers on seals and conduct aerial surveys concurrently, requiring considerably more personnel and money. One possible suggestion is to include year as a random effect within models for aerial survey counts such that, without specific knowledge of any particular year, the among-year variance is included in the modeled standard errors. Regardless of the specific approach, future estimates of Arctic seal abundance will require specific consideration of annual variability and changes in the timing of peak haul-out behavior when estimating trends, as one will not know if moderate differences in abundance estimates are attributable to changes in abundance or changes in haul-out behavior.

Predictions of absolute haul-out probability in this paper were somewhat different than those previously reported for these species, especially for bearded seals. For instance, Ver Hoef et al. (2014) and Conn et al. (2014) used haul-out correction factors with maximums of 0.66 for bearded seals, 0.62 for ribbon seals, and 0.54 for spotted seals, where these maximums corresponded to times near local solar noon in mid-late May. Results here suggest a haul-out probability on May 20 at local solar noon of 0.304 (95% CI: 0.258 – 0.354) for bearded seals across all age and sex classes, 0.715 (95% CI: 0.62 – 0.794) for adult male ribbon seals, 0.661 (95% CI: 0.576 – 0.738) for adult female ribbon seals, 0.742 (95% CI: 0.655 – 0.813) for adult male spotted seals, and 0.663 (95% CI: 0.574 – 0.742) for adult female spotted seals. Our estimates of haul-out probability reflect increased sample sizes in terms of number of individuals, inclusion of weather covariates, and improvements to the way data were prepared prior to analysis and should be the basis for any future estimates of seal abundance from aerial surveys in the Bering and Chukchi seas.

We focused this paper on haul-out behavior of bearded, ribbon, and spotted seals. Ringed seals are also present in the Bering and Chukchi seas but exhibit a unique complicating factor. Adult ringed seals build subnivean lairs under the snow on top of the sea ice, where they haul out and where females rear pups until conditions are good for basking (Frost et al., 2004). Thus, the wet-dry sensor on a bio-logger could indicate that an animal is hauled out, but if it is within a lair, it is not available to be detected during an aerial survey. We hope to address availability of ringed seals using data from satellite tags, replicate survey tracks, and auxiliary information about snow depth and timing of melt in a future study (see, e.g., Lindsay et al. (2021)).

Our analysis emphasizes the importance of accounting for seasonal and temporal variation in haul-out behavior, as well as associated weather covariates, when interpreting the number of seals counted in aerial surveys. The rapid acceleration of climate change in Arctic ecosystems is already occurring and is forecast to decrease the quantity and quality of suitable habitat for ice-associated seals that rely on sea-ice for a variety of activities. Improved estimates of seal population abundance from aerial surveys are needed to properly monitor the impacts of these changes on seal populations over time. Those monitoring surveys will need to be paired with continued investigation and assessment of seal haul-out behavior to allow credible predictions about the effects of climate disruptions on the abundance and distribution of Arctic seal populations.

## AUTHOR CONTRIBUTIONS

- **Josh M. London**: investigation, conceptualization, methodology, formal analysis, validation, software, writing: original draft, writing: review and editing, visualization, and data curation
- **Paul B. Conn**: conceptualization, methodology, formal analysis, software, validation, writing: original draft, writing: review and editing
- **Stacie K. Hardy**: investigation, data curation, methodology, validation, writing: review and editing
- **Erin L. Richmond**: data curation, investigation, methodology, validation, writing: review and editing
- **Jay M. Ver Hoef** : conceptualization, methodology, software, writing: review and editing
- **Michael F. Cameron**: investigation, project administration, writing: review and editing
- **Justin A. Crawford**: investigation, methodology, validation, data curation, writing: review and editing
- **Lori T. Quakenbush**: investigation, methodology, supervision, project administration, writing: review and editing
- **Andrew L. Von Duyke**: investigation, methodology, validation, data curation, writing: review and editing
- **Peter L. Boveng**: investigation, conceptualization, supervision, project administration, writing: review and editing

## DATA AVAILABILITY

This manuscript was developed as a reproducible research compendium. All data and code are available on GitHub (https://github.com/noaa-afsc/berchukseals-haulout) and major versions archived at Zenodo (https://doi.org/10.5281/zenodo.4638221). Original data sources for telemetry are archived at the United States Animal Telemetry Network, archived at Movebank, or associated with other published manuscripts (see supplemental material S1). Collated and cleaned data products needed to replicate the analysis along with the results of all model fits are also available and versioned as an R package on GitHub (https://github.com/noaa-afsc/ berchuksealsHauloutFits) and archived at Zenodo.

## ACKNOWLEDGMENTS

We recognize that the species and ecosystems we studied are within the ancestral and present-day environs of the Inpuiat and Yup’ik people who, through many uncredited contributions of traditional knowledge, provided early western naturalists and scientists with much of what gets described as the ‘basic biology’ of Arctic seals. The deployment of bio-logging devices used in this study were often done in collaboration with Alaska Native seal hunters and the approval of their communities. We would like to especially acknowledge the communities of Kotzebue, Koyuk, Nome, Nuiqsut, Scammon Bay, St. Michael, Utqiaġ vik, and Ulguniq (Wainwright) and the following individuals: James Adams, Jeff Barger, David Barr, Wendell Booth, Cyrus Harris, Nereus ‘Doc’ Harris, Grover Harris, Lee Harris, Tom Jones, Frank Garfield, Brenda Goodwin, Henry Goodwin, John Goodwin, Pearl Goodwin, Willie Goodwin, Brett Kirk, Noah Naylor, Virgil Naylor Jr., Virgil Naylor Sr., Dan Savetilik, Chuck Schaeffer, Ross Schaeffer, Allen Stone, and Randy Toshavik from Kotzebue; Merlin Henry from Koyuk; Tom Gray from Nome; Vernon Long and Richard Tukle from Nuiqsuit; Morgan Simon, River Simon, and Al Smith from Scammon Bay; Alex Niksik Jr. from St. Michael; Billy Adams, James Aiken, Tim Aiken, Howard Kittick, Gilbert Leavitt, Isaac Leavitt, J.R. Leavitt, and Joe Skin from Utqiaġ vik, Alaska; Mary Ellen Ahmaogak, Enoch Oktollik, Shawn Oktollik, Stacey Osborn, and Fred Rexford from Ulguniq.

We are grateful for the assistance in catching and sampling seals by Ryan Adam, James Bailey, Michelle Barbieri, John Bengtson, Gavin Brady, Vladamir Burkanov, Cynthia Christman, Sarah Coburn, Shawn Dahle, Rob Delong, Stacy DiRocco, Deb Fauquier, Shannon Fitzgerald, Kathy Frost, Scott Gende, Tracey Goldstein, Jeff Harris, Jason Herreman, Markus Horning, John Jansen, Shawn Johnson, Charles Littnan, Lloyd Lowry, Brett McClintock, Erin Moreland, Mark Nelson, Justin Olnes, Lorrie Rea, Bob Shears, Gay Sheffield, Brent Stewart, Dave Withrow, and Heather Ziel. We also appreciate the commitment to science and safety by all officers and crew of the NOAA ship *Oscar Dyson*, the NOAA ship *MacArthur II*, and the RV *Thomas G. Thompson*.

Telemetry data from the Alaska Department of Fish and Game (ADF&G) and the North Slope Borough Department of Wildlife Management (NSB) were important contributions to the findings presented here. Deployments in the western Bering Sea were done in collaboration with Russian colleagues and North Pacific Wildlife.

The findings and conclusions in the paper are those of the author(s) and do not necessarily represent the views of the National Marine Fisheries Service, NOAA. Any use of trade, product, or firm names does not imply an endorsement by the U.S. Government. Funding for this study was provided by the U.S. National Oceanic and Atmospheric Administration. The field work was conducted under the authority of Marine Mammal Protection Act Research Permits Nos. 782-1676, 782-1765, 15126, and 19309 issued by the National Marine Fisheries Service, and Letters of Assurance of Compliance with Animal Welfare Act regulations, Nos. A/NW 2010-3 and A/NW 2016-1 from the Alaska Fisheries Science Center/Northwest Fisheries Science Center Institutional Animal Care and Use Committee (IACUC). ). Funding to ADF&G for tagging seals was provided by the Bureau of Ocean Energy Management (No. M13PC0015 for work in Kotzebue, Alaska in 2009) and the Office of Naval Research (No. N00014-16-1-3019). ADF&G and NSB field work was covered by Research Permits Nos. 358-1585, 358-1787, 15324, and 20466 and by ADF&G IACUC permits Nos. 06-16, 09-21, 2014-03, 2015-25, 2016-23, 0027-2017-27, 0027-2018-29, 0027-2019-041.

## SUPPLEMENTAL MATERIAL

### 0.1 Additional Bio-logger Deployment Details

**Table S1.** The timing, location, and institutions responsible for the bio-logger deployments used in this study along with research permits and any associated publications.

| Institution | Year Deployed | Location | Publication(s) | Permits | No. Seals | Sex | Age Class |
| --- | --- | --- | --- | --- | --- | --- | --- |
| **Bearded seal** |  |  |  |  |  |  |  |
| ADFG | 2005 | Kotzebue Sound | Cameron et al 2018 | 358-1585 | 6 | M, F | subadult |
| ADFG | 2006 | Kotzebue Sound | Cameron et al 2018 | 358-1585 | 2 | F, M | subadult |
| ADFG | 2009 | Kotzebue Sound | Breed et al 2018 | 358-1787 | 4 | F, M | subadult |
| ADFG | 2014 | Norton Sound, Koyuk River | Olnes et al 2020 | 15324 | 2 | M | subadult |
| ADFG | 2014 | Norton Sound, Nome | Olnes et al 2020 | 15324 | 1 | M | subadult |
| ADFG | 2015 | Norton Sound, St. Michael | Olnes et al 2020 | 15324 | 1 | M | subadult |
| ADFG | 2016 | Elson Lagoon, Utqiagvik | Olnes et al 2020 | 15324 | 1 | F | subadult |
| ADFG | 2016 | Norton Sound, Koyuk River | Olnes et al 2020 | 15324 | 2 | F, M | subadult |
| ADFG | 2016 | Norton Sound, Nome | Olnes et al 2020 | 15324 | 1 | M | subadult |
| ADFG | 2016 | Norton Sound, St. Michael | Olnes et al 2020 | 15324 | 2 | M, F | subadult |

| **Institution** | **Year Deployed** | **Location** | **Publication(s)** | **Permits** | **No. Seals** | **Sex** | **Age Class** |
| --- | --- | --- | --- | --- | --- | --- | --- |
| ADFG | 2017 | Colville River, Nuiqsut | Olness et al 2020 | 15324 | 1 | F | subadult |
| ADFG | 2017 | Norton Sound, Koyuk River | Olness et al 2020 | 15324 | 1 | F | subadult |
| ADFG | 2017 | Norton Sound, Nome | Olness et al 2020 | 15324 | 1 | F | subadult |
| ADFG | 2019 | Dease Inlet, Utqiagvik | Olness et al 2021 | 20466 | 1 | M | adult |
| NMFS | 2005 | Kotzebue Sound |  | 358-1585 | 1 | F | subadult |
| NMFS | 2009 | Kotzebue Sound | McClintock et al 2017 | 782-1765 | 2 | M | subadult, adult |
| NMFS | 2011 | Kotzebue Sound | McClintock et al 2017 | 15126 | 3 | F, M | subadult |
| NMFS | 2012 | Kotzebue Sound | McClintock et al 2017 | 15126 | 1 | F | adult |
| NSB | 2012 | Elson Lagoon, Utqiagvik |  | 15324 | 1 | M | subadult |
| NSB | 2019 | Pittalugruaq Lake |  | 20466 | 1 | F | subadult |
| **Ribbon seal** |  |  |  |  |  |  |  |
| NMFS | 2005 | Ozemoy Gulf, Russia |  | 782-1765 | 9 | F, M | young of year, adult, subadult |
| NMFS | 2006 | Bering Sea |  | 782-1765 | 7 | M, F | adult, young of year |
| NMFS | 2007 | Bering Sea |  | 782-1765 | 28 | M, F | subadult, adult, young of year |

| **Institution** | **Year Deployed** | **Location** | **Publication(s)** | **Permits** | **No. Seals** | **Sex** | **Age Class** |
| --- | --- | --- | --- | --- | --- | --- | --- |
| NMFS | 2008 | Bering Sea |  | 782-1765 | 1 | M | subadult |
| NMFS | 2009 | Bering Sea |  | 782-1765 | 28 | F, M | adult, subadult, young of year |
| NMFS | 2010 | Bering Sea |  | 358-1787,<br>15126 | 17 | M, F | young of year, adult, subadult |
| NMFS | 2014 | Bering Sea |  | 15126 | 13 | M, F | subadult, adult, young of year |
| NMFS | 2016 | Bering Sea |  | 19309 | 7 | M, F | subadult, adult |
| **Spotted seal** |  |  |  |  |  |  |  |
| ADFG | 2005 | Kotzebue Sound | Von Duyke et al in prep | 358-1585 | 3 | F, M | subadult, adult |
| ADFG | 2016 | Dease Inlet, Utqiagvik | Von Duyke et al in prep | 15324 | 4 | M, F | adult |
| ADFG | 2017 | Colville River, Nuiqsut | Von Duyke et al in prep | 15324 | 1 | F | subadult |
| ADFG | 2017 | Scammon Bay | Von Duyke et al in prep | 15324 | 3 | F, M | subadult, adult |
| ADFG | 2018 | Dease Inlet, Utqiagvik | Von Duyke et al in prep | 20466 | 1 | F | subadult |
| ADFG | 2018 | Scammon Bay | Von Duyke et al in prep | 20466 | 1 | M | subadult |
| ADFG | 2019 | Dease Inlet, Utqiagvik | Von Duyke et al in prep | 20466 | 6 | M | adult, subadult |
| NMFS | 2006 | Bering Sea |  | 782-1676 | 5 | M, F | young of year, subadult |

| **Institution** | **Year Deployed** | **Location** | **Publication(s)** | **Permits** | **No. Seals** | **Sex** | **Age Class** |
| --- | --- | --- | --- | --- | --- | --- | --- |
| NMFS | 2007 | Bering Sea |  | 782-1676 | 12 | F, M | adult, young of year, subadult |
| NMFS | 2009 | Bering Sea |  | 358-1787 | 23 | F, M | adult, subadult, young of year |
| NMFS | 2010 | Bering Sea |  | 358-1787, 15126 | 8 | F, M | young of year, adult, subadult |
| NMFS | 2014 | Bering Sea |  | 15126 | 5 | M, F | young of year, adult |
| NMFS | 2016 | Bering Sea |  | 19309 | 6 | M, F | adult |
| NMFS | 2018 | Bering Sea |  | 19309 | 5 | F | adult |
| NPWC | 2009 | Kamchatka Peninsula |  | NA | 3 | F | adult |
| NSB | 2012 | Tiny Island | Von Duyke et al in prep | 15324 | 1 | F | adult |
| NSB | 2014 | Oarlock Island | Von Duyke et al in prep | 15324 | 6 | M, F | subadult, adult |
| NSB | 2014 | Seal Island | Von Duyke et al in prep | 15324 | 1 | M | subadult |
| NSB | 2015 | Oarlock Island | Von Duyke et al in prep | 15324 | 6 | M, F | subadult, adult |
| NSB | 2016 | Pittalugruaq Lake | Von Duyke et al in prep | 15324 | 3 | F | subadult |
| NSB | 2017 | Pittalugruaq Lake | Von Duyke et al in prep | 15324 | 1 | M | subadult |

**ADFG**=Alaska Department of Fish and Game; **NSB**=North Slope Borough; **NMFS**=NOAA National Marine Fisheries Service; **NPWC**=North Pacific Wildlife Consortium

### 0.2 Supplemental Figures Showing Confidence Intervals Associated with Predictions

The following series of figures (S1, S2, and S3) show the seasonal variability in predicted haul-out probability and the associated 95% confidence intervals for bearded, ribbon, and spotted seals. The predictions shown are based on the same data used in 5, 7, and 9 but selected for three local solar hours (07:00, 12:00, and 17:00) so the confidence intervals can also be shown and comparisons can be made.

**Figure S1.**
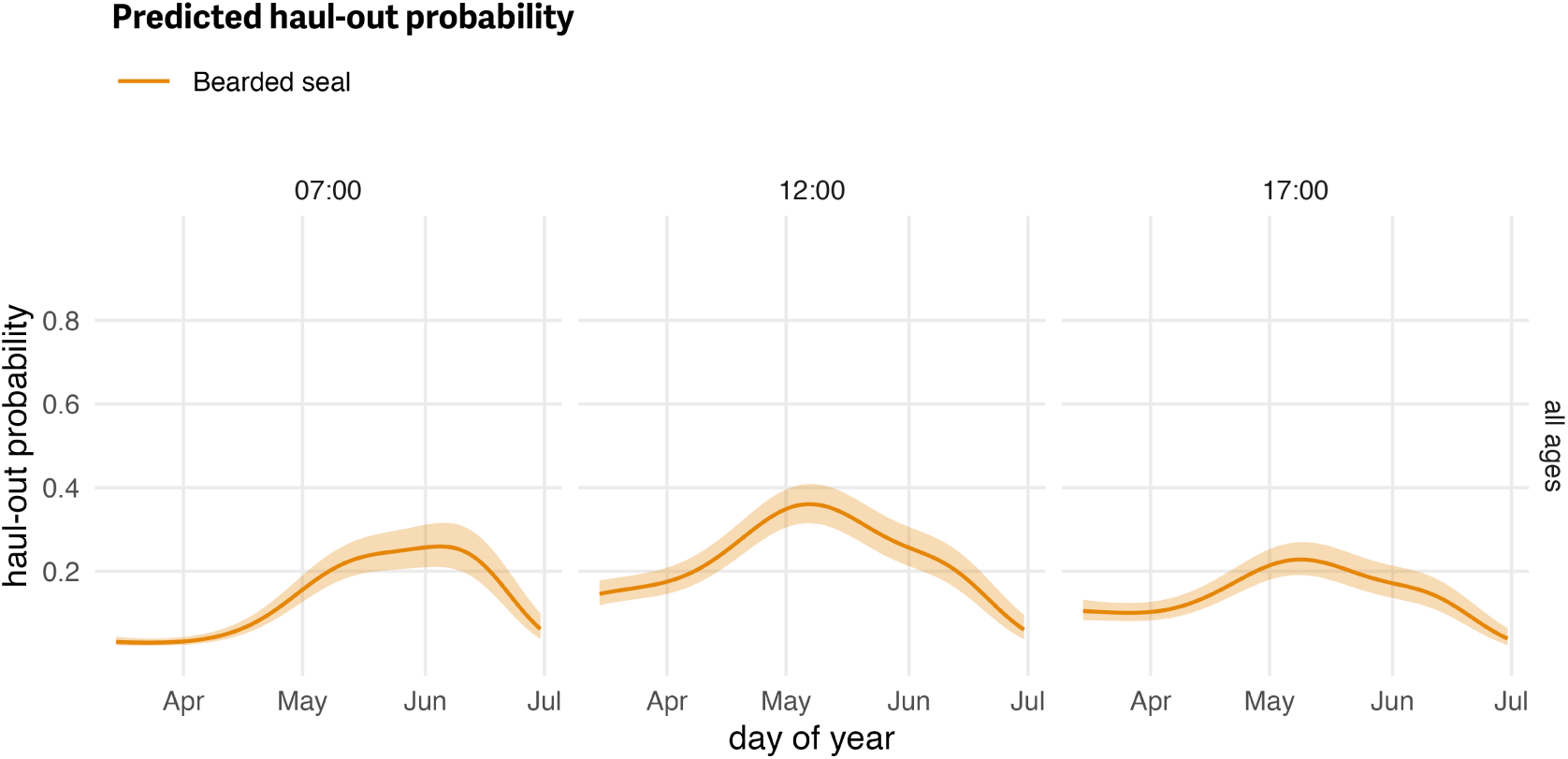
Seasonal variability in haul-out probability and the associated 95% confidence intervals (shaded area) for bearded seals. Model predictions are shown for three local solar hours (07:00, 12:00, and 17:00). Weather covariate values in the prediction were based on a simple generalized additive model for each weather covariate with smooth terms for day-of-year and solar hour to account for anticipated variability within a day over the season. Age and sex classes are combined into a single ‘all ages’ category.

**Figure S2.**
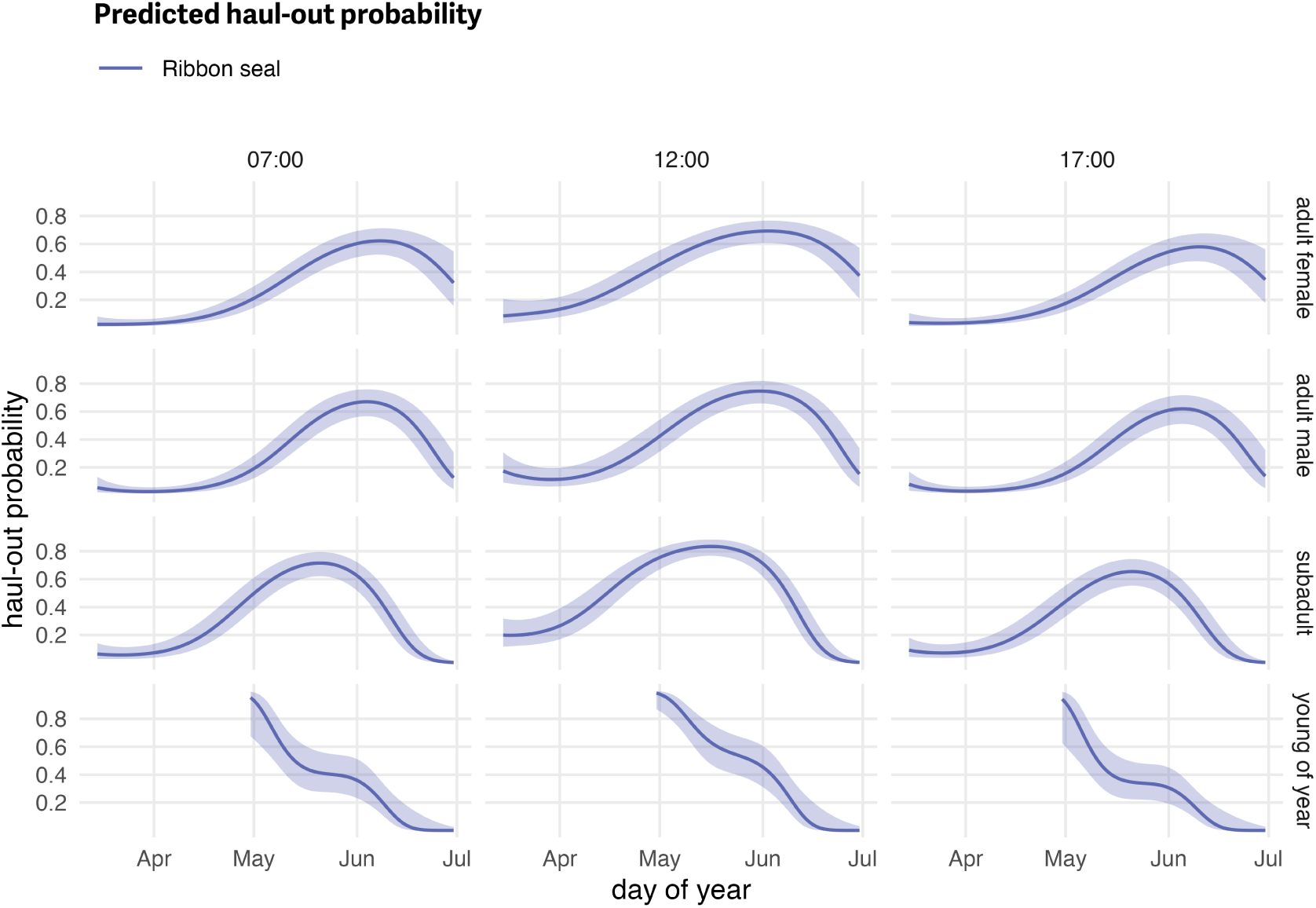
Seasonal variability in haul-out probability and the associated 95% confidence intervals (shaded area) for ribbon seals. Model predictions are shown for three local solar hours (07:00, 12:00, and 17:00). Weather covariate values in the prediction were based on a simple generalized additive model for each weather covariate with smooth terms for day-of-year and solar hour to account for anticipated variability within a day over the season. Age and sex classes are separated to allow comparisons.

**Figure S3.**
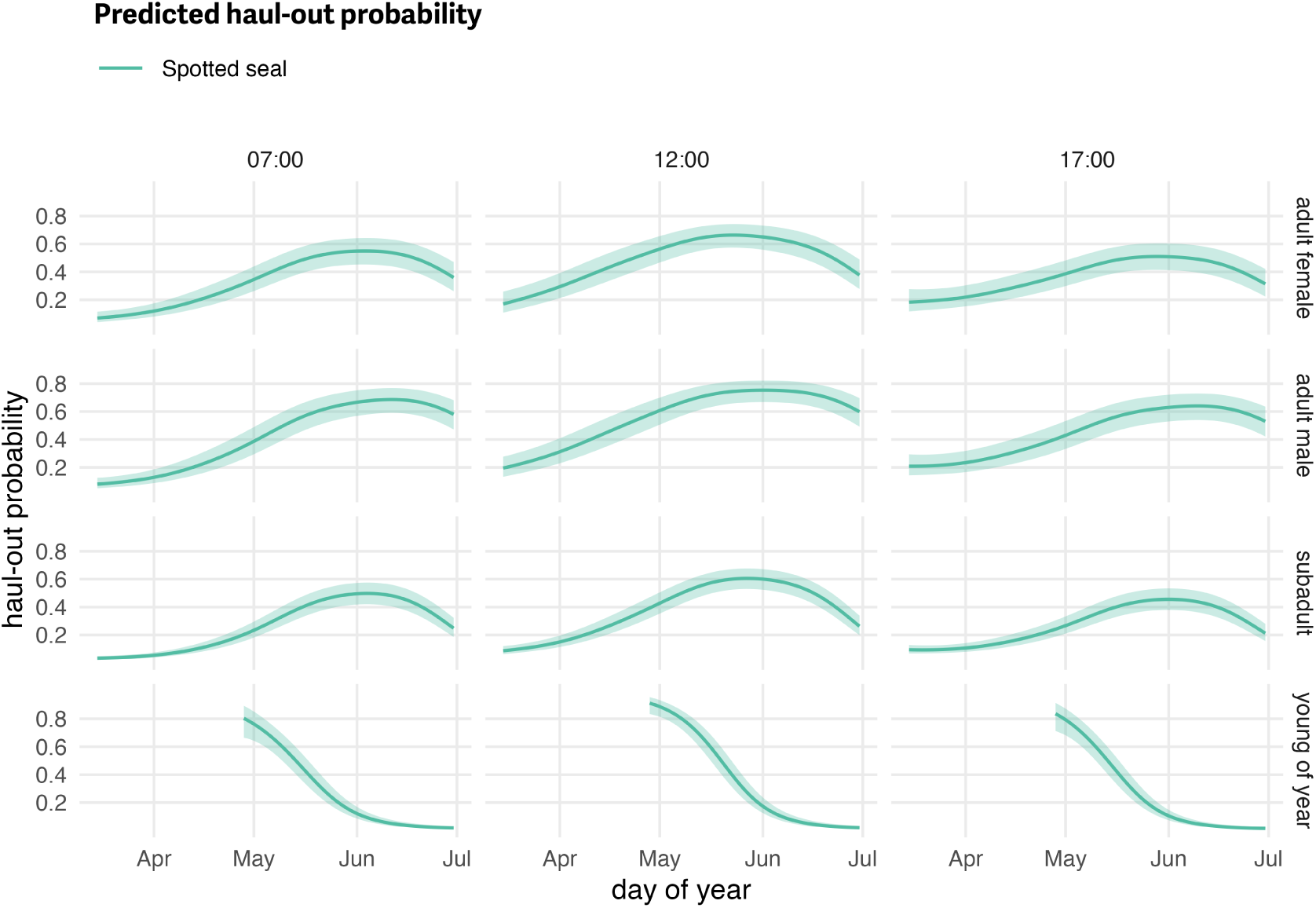
Seasonal variability in haul-out probability and the associated 95% confidence intervals (shaded area) for spotted seals. Model predictions are shown for three local solar hours (07:00, 12:00, and 17:00). Weather covariate values in the prediction were based on a simple generalized additive model for each weather covariate with smooth terms for day-of-year and solar hour to account for anticipated variability within a day over the season. Age and sex classes are separated to allow comparisons.

### 0.3 Exploring Insolation (Solar Radiation) as a Model Covariate

#### 0.3.1 Introduction

During the peer review process for this manuscript, Anthony Fischbach suggested the possibility of using predicted insolation (or solar radiation) values from the reanalysis model as a more direct and, potentially, more informative predictor of the daily haul-out cycle in seals compared to time of day. The notion being that seals are, likely, directly responding to changes in solar radiation throughout the day and not what time of day it is (i.e. seals don’t have human watches). Additionally, given the energetic benefits of increased solar radiation it could be more informative as we would expect seals might have a higher haul-out probability on sunnier days and for there to be geographic variability in haul-out behavior associated with geographical differences in insolation. This approach has an additional benefit of being more parsimonious compared to our use of the Fourier series or other approaches to represent hour-of-day in the model (e.g. 24 factors for each hour).

Because of these reasons, we considered and explored this possibility for our model and the analysis presented in this manuscript. A key drawback to reliance on solar radiation, in our minds, is that we would lose insight regarding potential diel patterns – solar radiation does not differentiate between dusk or dawn. Bi-modal patterns have been previously observed in ringed seals and our results in this analysis show some indication of increased haul-out probability during dawn compared to dusk periods for bearded seals and some age and sex classes for ribbon and spotted seals. For other phocid species, increased haul-out probability before solar noon or after solar noon has been observed. Importantly, understanding these relationships between haul-out probability and hour-of-day can have important ramifications on aerial survey study design – a key focus of this paper.

Another hesitation we had was that solar radiation estimates from reanalysis models have not been previously used as a model covariate within a published study of pinniped haul-out behavior. Thus, for this analysis, we chose to keep our original approach and rely on the Fourier series to capture any hour-of-day effects.

That said, we think the idea of solar radiation as a model covariate in pinniped haul-out models is intriguing and worth further exploration. The current availability and increased accessibility to detailed climate reanalysis products that include solar radiation is exciting and we encourage future, more detailed exploration of this as a component in pinniped haul-out analysis. To provide some inspiration, we present some initial efforts and examples for comparison.

### 0.3.2 Methods

In this manuscript, we rely on the NARR reanalysis model as the source for our weather covariates. However, since our initiation of this analysis, the ERA5 reanalysis model (https://doi.org/10.24381/cds.adbb2d47) has become one of the go-to standards for global climate reanalysis and provides an increased temporal resolution to hourly (compared to the 3-hour resolution of NARR). The global coverage of ERA5 provides additional flexibility in that the area of interest is not limited to North America. The ERA5 model provides a number of solar radiation parameters and it was important to evaluate and understand each of these estimates in order to select the one that was likely most relevant to seals. Here, we used the ‘surface short-wave (solar) radiation downwards’ parameter. This parameter is described as *“the amount of solar radiation (also known as shortwave radiation) that reaches a horizontal plane at the surface of the Earth and comprises both direct and diffuse solar radiation. To a reasonably good approximation, this parameter is the model equivalent of what would be measured by a pyranometer (an instrument used for measuring solar radiation) at the surface”* (https://codes.ecmwf.int/grib/param-db/?id=169). Thus, this is the value which most closely represents the amount of solar radiation likely felt by a seal hauled out of the water.

ERA5 data is available via the Copernicus climate data store API which can be queried with the CDS-API Python package (https://cds.climate.copernicus.eu/api-how-to). The R code provided here documents the download of the *surface_solar_radiation_downwards* parameter for our study area of interest and years of interest. The `reticulate` R package (https://CRAN.R-project.org/package=reticulate) allowed interaction with Python. Additionally, note, extra steps are required to download data on either side of the 180 anti-meridian.

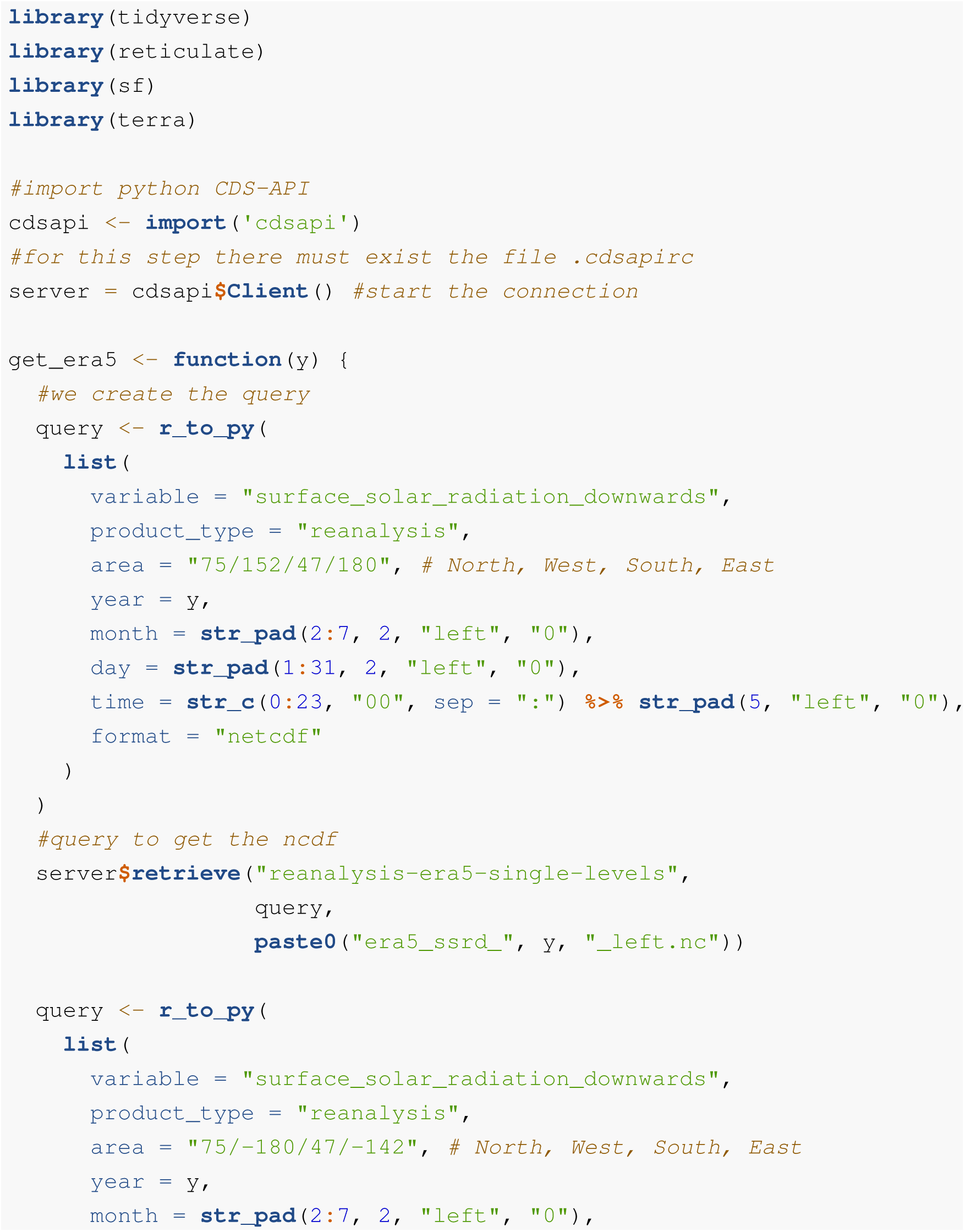
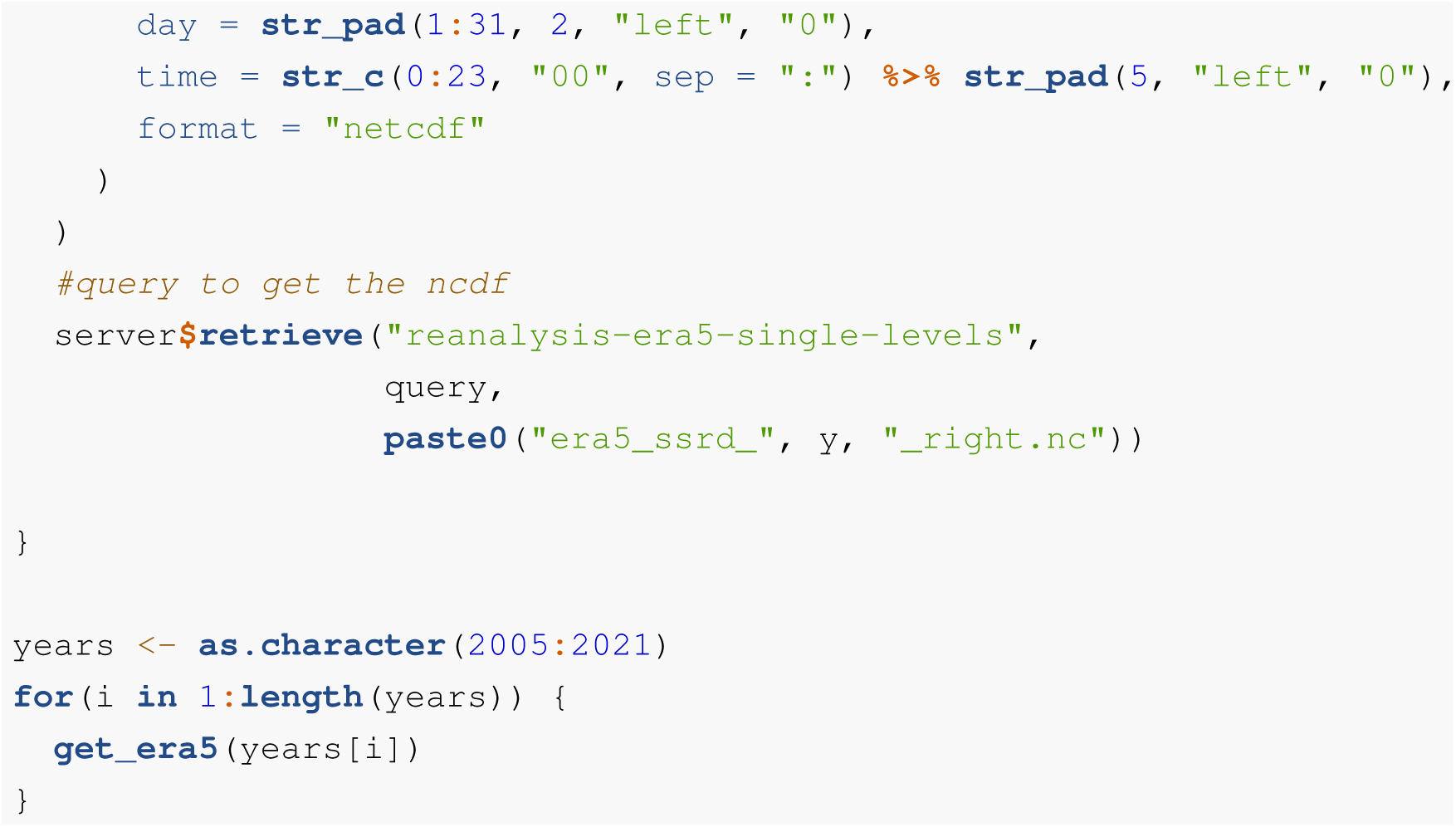

To explore performance of our solar radiation parameter within a haul-out model we replaced the various Fourier series parameters in our model from the manuscript with the ERA5 *surface solar radiation downwards* (`era_ssrd_watts`) parameter. As with other reanalysis values (from NARR) in the manuscript, the `era-ssrd-watts` values are matched in time and space to the seal haul-out observation data; we use the full hourly temporal resolution from ERA5. The glmmLDTS framework used in the paper does not allow for model comparisons with AIC because of the reliance on pseudo-likelihood. The bam() function within the `mgcv` package provides a quick model fitting option that also allowed us to do some model comparison with AIC. This approach was sufficient for the general demonstration and exploration purposes here but future research should consider a range of model fitting frameworks and approaches that might be more appropriate.

The model specification below was used to specify an `mgcv::bam()` model that matched the formula used in the manuscript for ribbon seals. The `s(speno, bs = "re")` term is the smooth term for the random effect. All other predictors were the same.

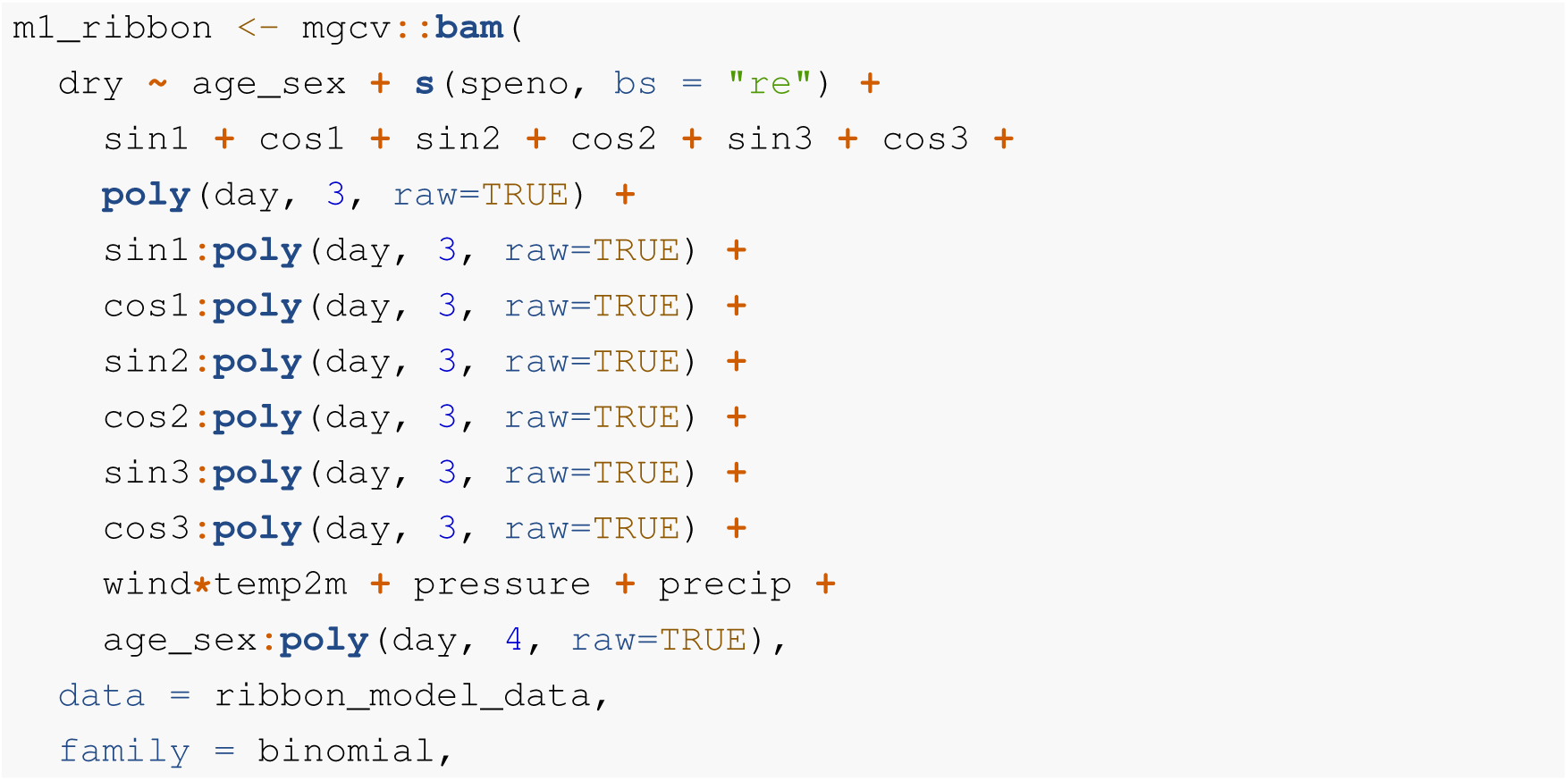
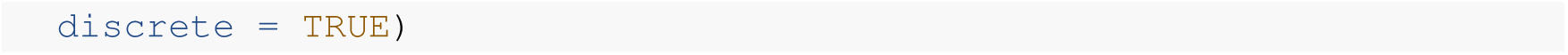

Note, the specification for *m1_ribbon* here does not include any AR1 structure for temporal autocorrelation. To include this, we needed to provide a value for 𝜌 (or *rho*). We examined the autocorrelation within the model and used the lag-1 value for 𝜌 .The value for lag-1 autocorrelation was 0.8082 which is rather high but not surprising. We then updated our model specification with a value for 𝜌 as well as the `A1.start` argument which specifies (as either **TRUE** or **FALSE**) the start point of each block.

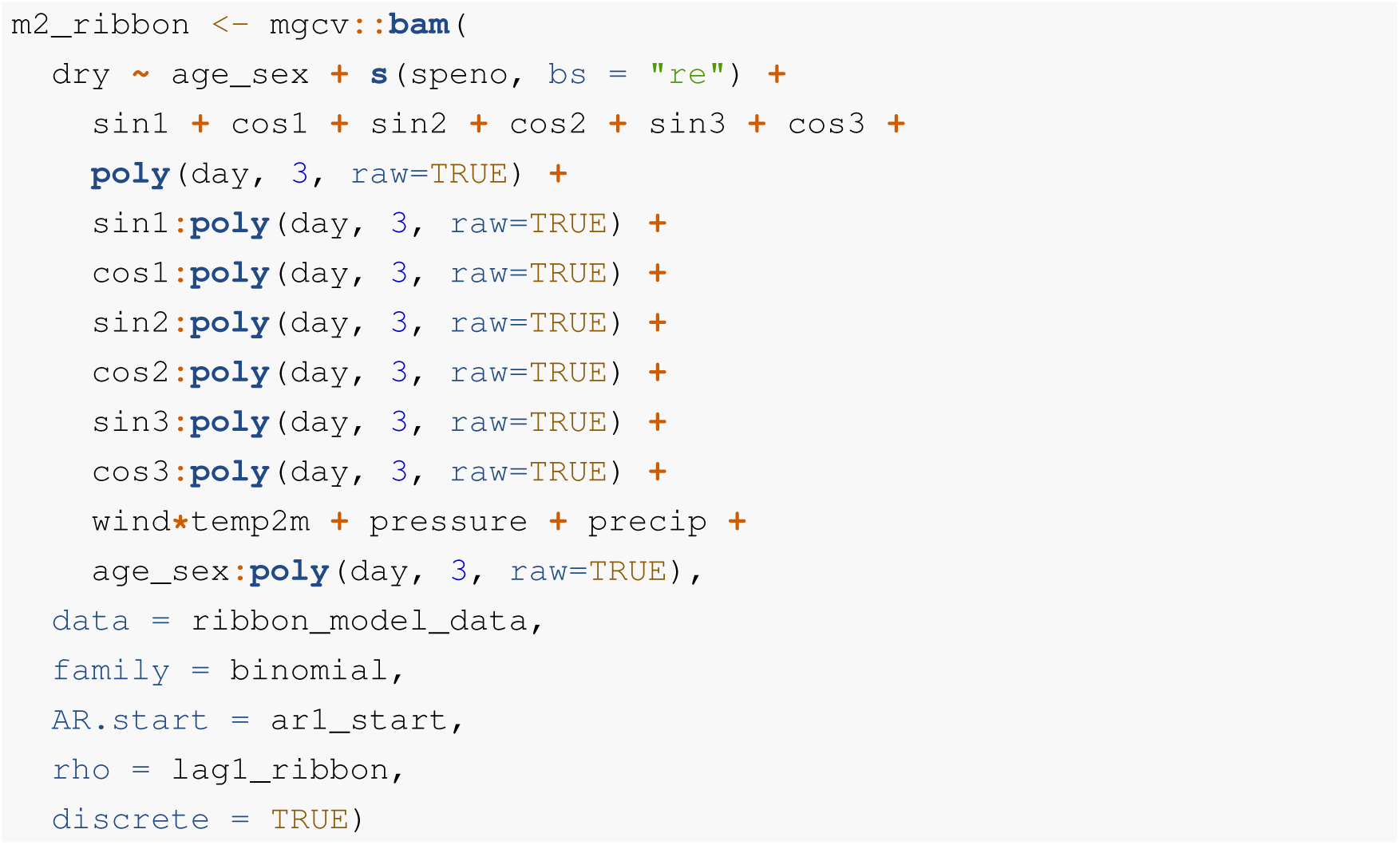

The model specification for exploring the use of solar radiation was specified similarly but without all of the Fourier series parameters and interactions.

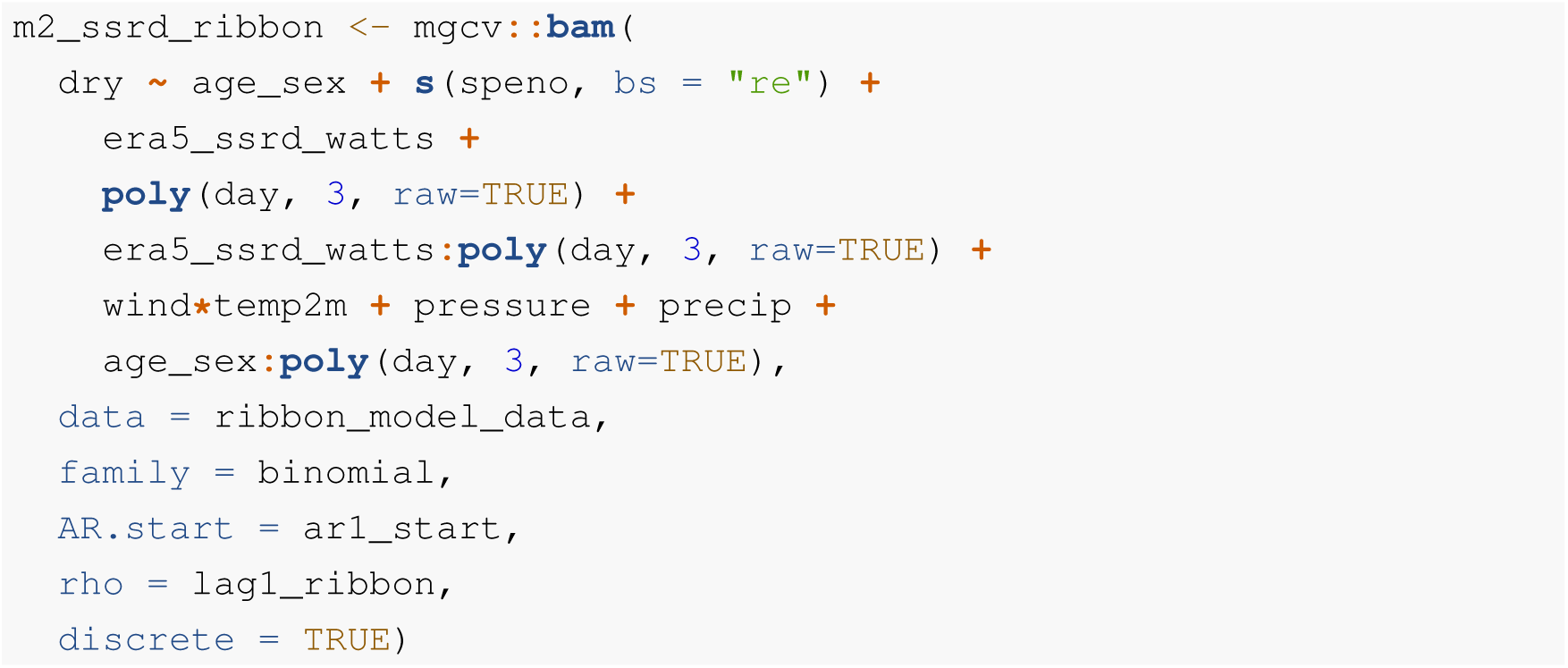

The two models were compared with AIC to evaluate whether the reduction in degrees of freedom with fewer terms in the solar radiation model was matched with improved explanatory power in the model fit. While the model and code specified above is for ribbon seals, the same approach was repeated for bearded and spotted seals.

**Figure S4.**
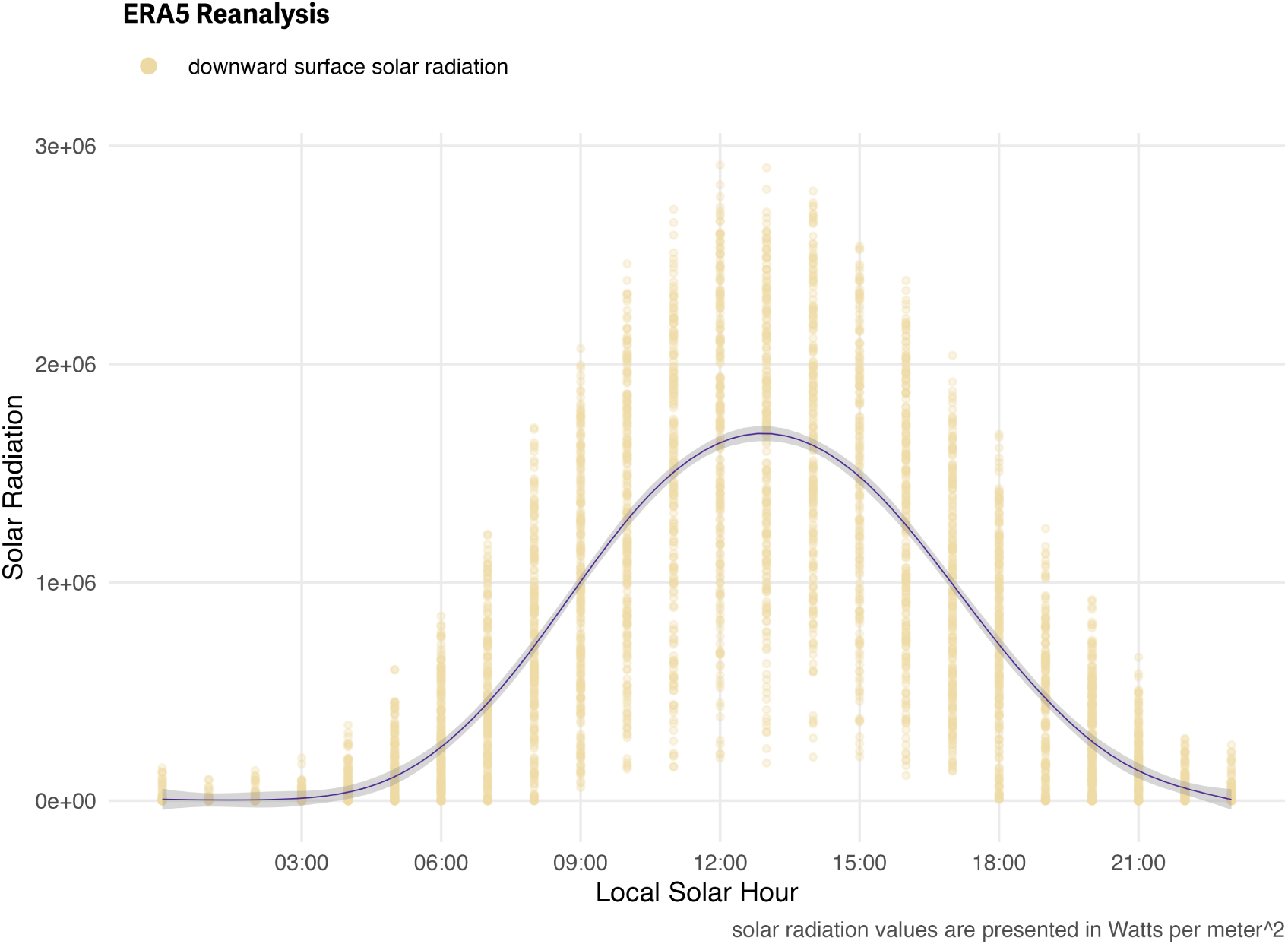
Diel Pattern of Solar Radiation Values from ERA5 Reanalysis. Downward surface solar radiation estimates from the ERA5 climate reanalysis for 5000 random points within the study area between 2005 and 2021. Solar radiation values are presented in Watts per square-meter and the smoothed line highlights the strong diel pattern.

A similar approach to that presented in this manuscript for prediction was employed with solar radiation values in lieu of hour of day. For prediction values, quantiles (5% increments) of the observed range of ERA5 solar radiation values were used with 100% representing the maximum observed solar radiation value. This allowed similar data visualizations and easier comparisons to those predictions in the manuscript that include hour of day.

#### 0.3.3 Results

To evaluate whether the solar radiation parameter matched our expectations and compared well with hour of the day, we visualized the variability of the `era5_ssrd` values within our study area as they relate to hour of the day (S4). The unimodal distribution is centered around the middle of the solar day with peak solar radiation coinciding with 13:00 local solar. This suggests solar radiation could be an informative covariate for capturing unimodal diel patterns in haul-out behavior.

The bearded seal model matching the specification from the manuscript resulted in 126.13 degrees of freedom and an AIC value of -7428.929. The model with solar radiation resulted in 39.829 degrees of freedom and an AIC value of -6874.298. The ribbon seal model matching the specification from the manuscript resulted in 131.478 degrees of freedom and an AIC value of -16372.29. The model with solar radiation resulted in 114.72 degrees of freedom and an AIC value of -15223.346. The spotted seal model matching the specification from the manuscript resulted in 125.54 degrees of freedom and an AIC value of -23482.051. The model with solar radiation resulted in 108.985 degrees of freedom and an AIC value of -22917.167. Despite the additional terms, the models with the Fourier series representation of hour of day resulted in a lower AIC value and were still preferred models for each of the species.

Predictions from the model fits and visualization of those predictions were produced for each species but, here, we only present visualizations from ribbon seals as an example (Figure S5 and Figure S6). Similar seasonal patterns previously observed were still apparent with subadults hauling out earlier in the season followed by adult males and, then, adult females. The observed relationship with hour of day and the centering of peak haul-out probability around solar noon was reflected in these predictions as a one-sided distribution with maximum solar radiation having the highest haul-out probability and minimal solar radiation the least. The seasonal distribution of haul-out probability along with 95% confidence intervals also provided comparable insights (see figures S2 and S6). That said, subtle differences in the shape and extent of confidence limits were present.

#### 0.3.4 Discussion

Solar radiation has potential as an informative covariate in pinniped haul-out models that can be directly linked to seal physiology and expected behavioral changes. The ERA5’s *surface solar radiation downwards* values aligned with hour of day and maximum values occurred at or just after local solar noon. This highlighted the informative potential for this approach. However, despite an overall reduction in the total number of parameters and degrees of freedom, AIC comparison still favored the models for each species that included hour of day as a Fourier series.

This analysis was not intended to be a full comparison – we simply want to demonstrate the potential and inspire further investigation – but, there are three possibilities that might explain the preference for hour of day. First, there are a broad range of solar radiation values represented for each hour of the day. Cloud cover, fog, and precipitation all reduce downward solar radiation at the surface and we might expect this to impact haul-out probability. However, the photoperiod and the timing of sunrise and sunset are not impacted by weather and seals may be responding to these signals more than the amount of solar radiation. Additionally, this study spans a range of physiological cycles and energetic needs and higher solar radiation may not be a consistent driving influence on seals. Increased energy from the sun may be important during molt but less so during pupping and breeding periods. Second, the timing and duration of haul-out behavior may also be influenced by diel patterns in weather (e.g. lower winds in the morning) or ecosystem dynamics (e.g. prey availability) that lead to a skewness in the distribution of haul-out behavior that wouldn’t be reliably captured by solar radiation values. Third, this effort is only an initial effort to explore the use of solar radiation in pinniped haul-out models. A more in depth and rigorous exploration of this topic might discover an approach that results in a more parsimonious and preferred model formulation.

Again, we want to acknowledge Anthony Fischbach for the suggestion during the peer review process. We think this is an excellent example of the peer review process working to improve the quality of our manuscript and advance the scientific process. We hope others will take our example and expand on it within future analyses.

**Figure S5.**
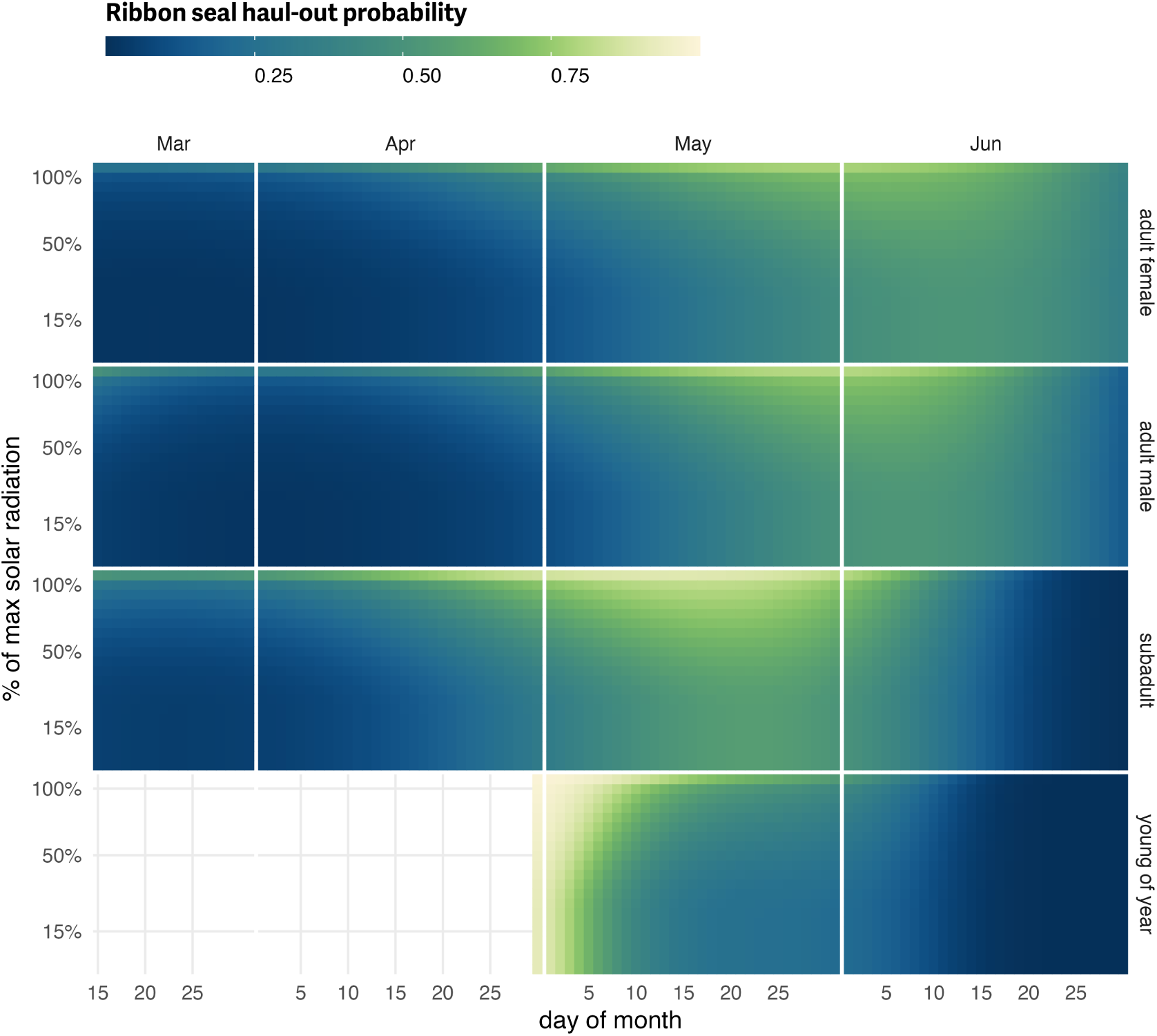
Solar radiation as a predictor of ribbon seal haul-out probability. Predicted haul-out probability of ribbon seals from 15 March to 30 June for each age and sex class used in the model. In this model, solar radiation was used in lieu of hour of day. The apparent seasonal progression with subadults hauling out earlier in the season followed by adult males and, then, adult females is still notable although maybe not as clear. Predictions for young of the year still show their transition from newly weaned pups resting on the ice to more in-water activities. The overall pattern is in agreement with a one-sided view of Figure 7 where maximum solar radiation is equivalent to local solar noon.

**Figure S6.**
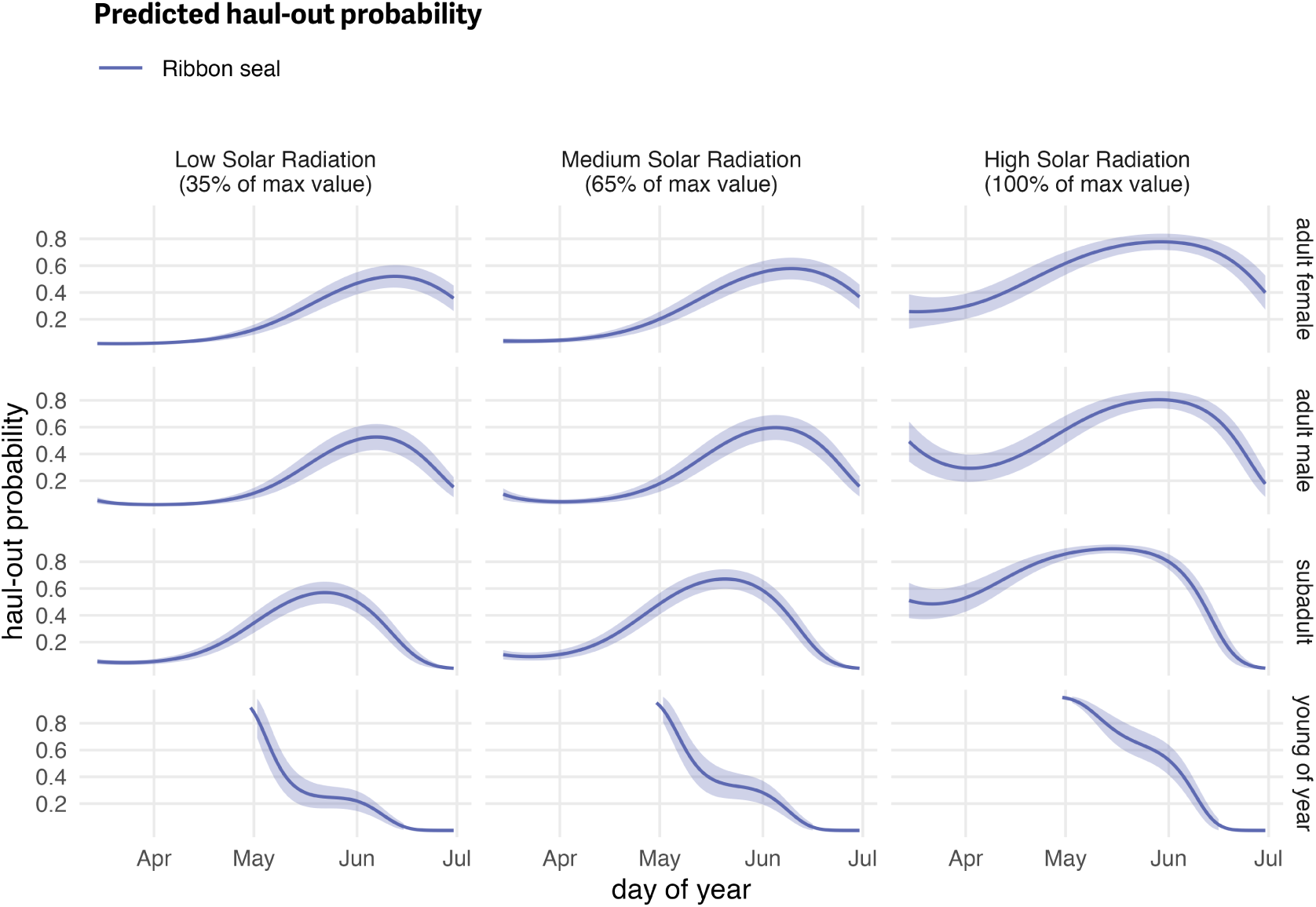
Solar radiation as a predictor of ribbon seal haul-out probability (with uncertainty). Seasonal variability in haul-out probability and the associated 95% confidence intervals (shaded area) for ribbon seals. In this model predictions are shown for low, medium, and high values of solar radiation (as percentages of the maximum value observed) in lieu of local solar hour. There’s general agreement in the overall seasonal patterns between the two approaches but sublte differences in shape and extent of the confidence limits were present (see Figure S2 for comparisons).

